# Displacement of PKA catalytic subunit from AKAP signaling islands drives pathology in Cushing’s syndrome

**DOI:** 10.1101/2021.10.18.464848

**Authors:** Mitchell H. Omar, Dominic P. Byrne, Kiana N. Jones, Tyler M. Lakey, Kerrie B. Collins, Kyung-Soon Lee, Leonard A. Daly, Katherine A. Forbush, Ho-Tak Lau, Martin Golkowski, G. Stanley McKnight, David T. Breault, Anne-Marie Lefrançois-Martinez, Antoine Martinez, Claire E. Eyers, Geoffrey S. Baird, Shao-En Ong, F. Donelson Smith, Patrick A. Eyers, John D. Scott

## Abstract

Mutations in the catalytic subunit of protein kinase A (PKAc) drive the stress hormone disorder adrenal Cushing’s syndrome. Here we define mechanisms of action for the PKAc-L205R and W196R variants. Both Cushing’s mutants are excluded from A kinase anchoring protein (AKAP) signaling islands and consequently diffuse throughout the cell. Kinase-dead experiments show that PKA activity is required for cortisol hypersecretion. However, kinase activation is not sufficient, as only cAMP analog drugs that displace native PKAc from AKAPs enhance cortisol release. Rescue experiments that incorporate mutant PKAc into AKAP signaling islands abolish cortisol overproduction, indicating that kinase anchoring restores normal endocrine function. Phosphoproteomics show that PKAc-L205R and W196R engage different mitogenic signaling pathways. ERK activity is elevated in adrenal-specific PKAc-W196R knock-in mice. Conversely, PKAc-L205R attenuates Hippo signaling, thereby upregulating the YAP/TAZ transcriptional co-activators. Thus, aberrant localization of each Cushing’s variant promotes the transmission of a distinct downstream pathogenic signal.

## Introduction

Cushing’s syndrome is a disease of chronically elevated levels of stress hormone. Patients of this endocrine disorder classically present with weight gain, a moon face, chronic fatigue, hypertension, and depression, and are most often diagnosed between the ages of 20 and 50 (Lacroix et al., 2015). At the molecular level, Cushing’s syndrome is predominately a consequence of defective cAMP signaling (Lacroix et al., 2015). This can proceed via elevated adrenocorticotropic hormone (ACTH), or result from genetic lesions in cAMP signaling elements that operate within the adrenal gland zona fasciculata, a mid-zone of the adrenal cortex (Hernandez-Ramirez and Stratakis, 2018; Rosenberg et al., 2002). Whole exome sequencing of adrenal tissue from patients has identified mutations and insertions in protein kinase A catalytic subunits (PKAc) that are drivers of this endocrine disorder (Beuschlein et al., 2014; Cao et al., 2014; Di Dalmazi et al., 2014; Espiard et al., 2018; Goh et al., 2014; Ronchi et al., 2016; Sato et al., 2014). Most Cushing’s PKAc mutations cluster where the kinase interfaces with its regulatory subunits (R) (Bathon et al., 2019). This protein-protein interaction is not only necessary for autoinhibition of kinase activity, but also directs the subcellular targeting of PKA holoenzymes through association with A kinase anchoring proteins (AKAPs) (Omar and Scott, 2020; Taylor et al., 2012).

A kinase anchoring proteins (AKAPs) are a growing family of scaffolding proteins that organize and direct subcellular signaling events (Fig 1A) (Omar and Scott, 2020; Scott et al., 2013; Taskén and Aandahl, 2004). Three characteristic properties of AKAPs mediate their function. First, enzyme binding sites on each AKAP permit the assembly of distinct combinations of signaling molecules (Klauck et al., 1996). Second, differential utilization of subcellular targeting motifs constrains each cohort of anchored enzymes within defined intracellular environments such as the plasma membrane, centrosomes, mitochondria, and other organelles (Wong and Scott, 2004). Third, anchoring of PKA within cAMP nanodomains creates discrete “signaling islands” where this kinase is sequestered near its preferred substrates (Fig 1A) (Bock et al., 2021; Musheshe et al., 2018; Smith et al., 2017). The defining molecular characteristic of AKAPs is a conserved amphipathic helix that binds to the regulatory(R) subunits of PKA with high affinity (Burgers et al., 2015; Carr et al., 1991). Since the regulatory subunits constrain PKAc, they provide a means to localize kinase activity (Smith et al., 2017; Smith et al., 2013). Moreover, type I and type II PKA holoenzymes can be differentially recruited to type I- or type II-selective AKAPs (Means et al., 2011; Taylor et al., 2012). Since most cells typically express 15-20 different AKAPs, they create an archipelago of signaling islands that can simultaneously propagate individual PKA phosphorylation events efficiently and with minimal pathway crosstalk (Langeberg and Scott, 2015). The importance of this local signaling mechanism is underscored by evidence that loss of AKAP-mediated enzyme targeting disrupts essential physiological processes such as learning and memory, cardiac function, and insulin release (Hinke et al., 2012; Lygren et al., 2007; Nieves-Cintron et al., 2016; Sanderson et al., 2016; Tunquist et al., 2008).

**Figure 1.**
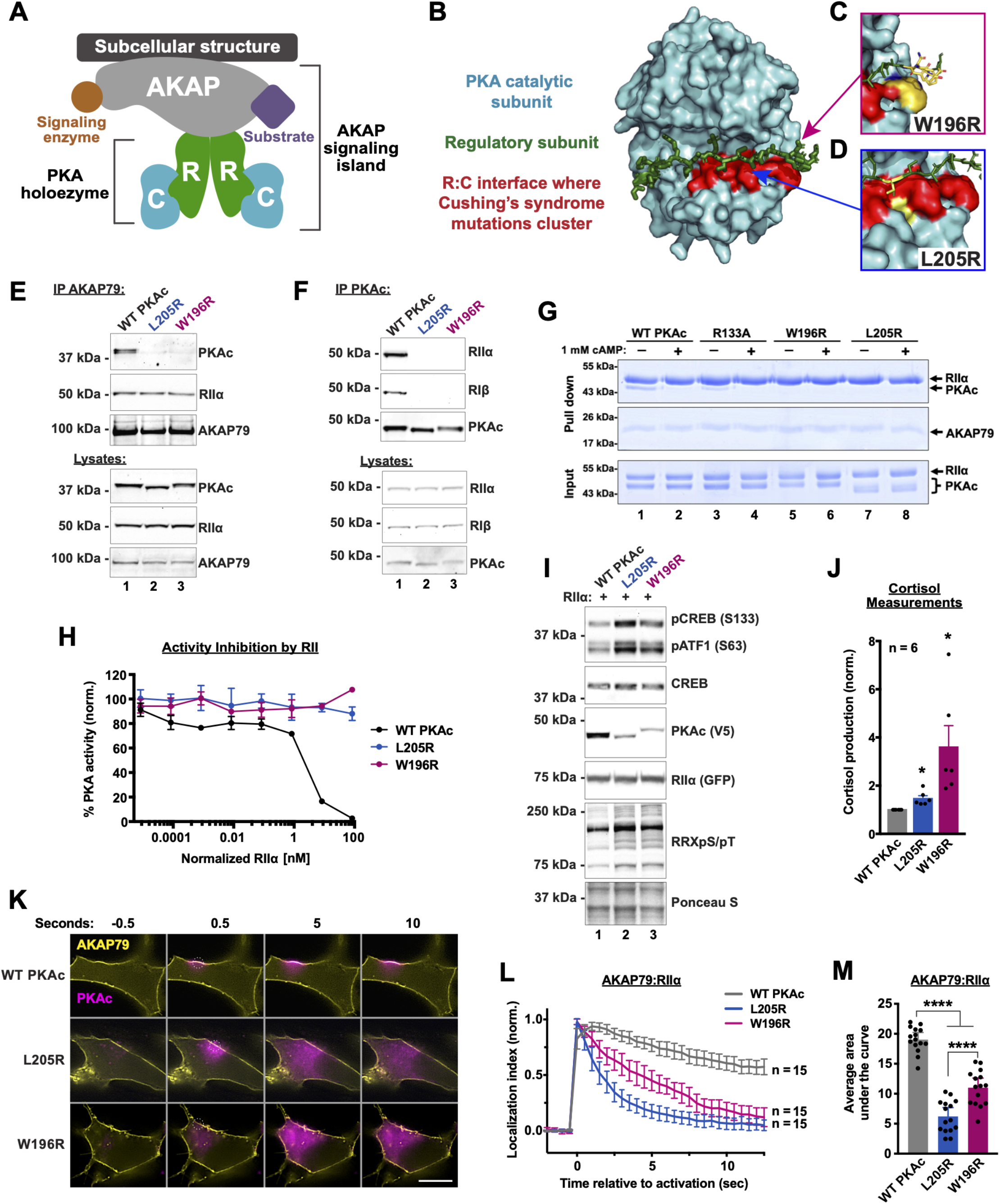
Cushing’s mutants are excluded from AKAP signaling islands. A) Model of AKAP signaling island. Regulatory subunits, R; Catalytic subunits, C. B) Structure of PKAc bound to the inhibitory region of RIIβ (green). Red region highlights residues where mutations have been identified in adrenal Cushing’s syndrome. The W196R (C) and L205R (D) mutations fall within this region. E) Immunoprecipitation of AKAP79 from H295R adrenal cells. WT, but not mutant, PKAc coprecipitates with AKAP signaling islands. F) Immunoprecipitation of PKAc from H295R cells. RI and RII coprecipitate with WT, but neither mutant, kinase. G) AKAP79(297-427)-PKA complex formation in vitro (purified recombinant proteins) in the presence/absence of cAMP. H) Changes in PKA dependent phosphorylation of a fluorescently labeled peptide substrate in the presence of increasing concentrations of RIIα regulatory subunit. Data shown is mean and standard deviation of 4 experiments. I) Immunoblots of CRISPR PKAcα-/- U2OS cells expressing RII-GFP along with either WT or mutant PKAc-V5. J) Cortisol measurements from H295R cells 72 h after lentiviral infection with PKAc variants. *p ≤ 0.05 determined by posthoc students t-tests after 1-way ANOVA. n = 6. K) Photoactivation timecourses in H295R cells. AKAP79-YFP, RII-iRFP, and PKAc tagged with photoactivatable mCherry were expressed. Scale bar = 10 μm. L) Amalgamated quantitation of K. Over time, the majority of WT PKAc (gray) remains localized to AKAP79 while L205R (blue) and W196R (red) diffuse away. Localization index = fluorescence intensity at the site of activation divided by intensity at off target sites. Data represents 3 experimental replicates. M) Integration of values in L. ****p ≤ 0.0001, corrected for multiple comparisons with Dunnett method after 1-way ANOVA.

PKAc is a promiscuous kinase that phosphorylates hundreds of cellular substrates (Shabb, 2001). Accordingly, mutations, insertions, and fusions of the *PRKAC* and *PRKAR* families of genes are linked to a growing number of diseases (Ma et al., 2019; Palencia-Campos et al., 2020; Stratakis, 2018). In this report, we focus on the mechanism of two mutations in PKAc that occur in patients suffering from adrenal (adrenocorticotropic hormone-independent) Cushing’s syndrome. Both PKAc mutations occur at the interface that binds PKA regulatory subunits (Fig 1B-D). The most prevalent mutant identified is L205R, which is found in ~45% of Cushing’s patients, fails to bind the R subunit of PKA, and is constitutively active (Bathon et al., 2019; Calebiro et al., 2014). Seven other PKAc mutations including the W196R variant have also been discovered in adenomas of Cushing’s patients (Bathon et al., 2019; Ronchi et al., 2016). Herein, we report similarities and differences in the mechanisms of action of PKAc-L205R and PKAc-W196R. Both mutations favor the intracellular diffusion of PKAc and boost stress hormone secretion from adrenal cells. Yet, proximity phosphoproteomics and functional studies reveal that PKAc-L205R and PKAc-W196R engage distinct mitogenic signaling cascades to exert their pathogenic mechanisms of action.

## Results

We first assessed to what extent the Cushing’s syndrome mutations in PKAc disrupt association with AKAP signaling islands in human adrenal NCI-H295R cells. Immunoprecipitation of a tagged AKAP79 successfully pulled down regulatory subunits and wild type (WT) PKAc (Fig 1E). In contrast, PKAc-L205R and PKAc-W196R were not detected when AKAP79 immunoprecipitation was performed in cells expressing the Cushing’s variants (Fig 1E, lanes 2 & 3). We next tested association between regulatory subunits and each PKAc form. Immunoprecipitation of WT PKAc successfully pulled down both type I and type II regulatory subunits (Fig 1F, lane 1). Neither L205R nor W196R co-precipitated the regulatory subunits (Fig 1F, lanes 2 & 3). Proteins purified from bacteria were subjected to in vitro experiments to more rigorously evaluate effects of L205R or W196R substitutions on protein-protein interactions within AKAP signaling islands. Upon pulldown of AKAP79, RII was detected in all conditions (Fig 1G, top panel). WT PKAc and PKAc-R133A, a variant known to be deficient in PKI binding, retained the ability to bind RII (Fig 1G, lanes 1 & 3) (Wen and Taylor, 1994). Control experiments confirmed that incubation with supraphysiologic levels of cAMP disrupted all PKAc binding (Fig 1G, lanes 2 & 4). In contrast, RII failed to pull down L205R or W196R variants of PKAc with or without excess cAMP (Fig 1G, lanes 5-8). Loading controls confirm that equivalent levels of AKAP79 and the PKA RII and C subunits were added in each experimental condition (Fig 1G, mid and bottom panels).

Regulatory subunits autoinhibit PKAc activity (Builder et al., 1980). Accordingly, titration of RII causes a precipitous decrease in WT PKA activity between RII concentrations of 0.83 nM and 8.3 nM (Fig 1H, black trace). In contrast, neither Cushing’s variant PKAc-L205R (blue) nor PKAc-W196R (red) is autoinhibited by RII (Fig 1H; supplemental fig 1A). These experiments support evidence that the Cushing’s syndrome mutants are tonically active inside cells (Beuschlein et al., 2014; Gibson and Taylor, 1997; Orellana and McKnight, 1992). To test this further, we examined the phosphorylation of the canonical PKA substrates CREB and ATF1 in a genetic background that was depleted of native kinase. CRISPR-Cas9 gene editing was used to remove PKAcα, the predominant kinase isoform in U2OS cells (Smith et al., 2017). Rescue experiments were performed by reintroducing PKAc-L205R or PKAc-W196R. Immunoblot detection of phospho-CREB and phospho-ATF1 served as an index of kinase activity (Fig 1I). Under basal conditions the WT kinase minimally phosphorylated either substrate (Fig 1I, lane 1). In contrast, robust phosphorylation was detected in conditions expressing either Cushing’s mutant (Fig 1I, lanes 2 & 3). In addition, each Cushing’s mutation displays increased phosphorylation of PKA substrates, as detected with an antibody to phosphorylated PKA target sequence RRXS/T (Fig 1I, fifth panel). Next, we measured human stress hormone (cortisol) release from H295R cells. Control experiments established baseline cortisol release from H295R adrenal cells expressing WT PKAc for 72 h (Fig 1J, gray). Cortisol release was elevated in conditions expressing PKAc-L205R (Fig 1J, blue). Interestingly, expression of PKAc-W196R had the most robust affect showing over three-fold production of stress hormone as compared to expression of the WT kinase (Fig 1J, red). Collectively, these structural, biochemical, and cell-based data suggest both L205R and W196R mutations disrupt inhibition of PKAc by its regulatory subunits and displace the kinase from AKAP signaling islands.

To visualize how Cushing’s mutations impact localization and mobility, we used live-cell photoactivation microscopy. Upon activation of a 2 μM diameter circle on the membrane of H295R cells, WT PKAc tagged with photoactivatable mCherry maintained its localization relative to fluorescently tagged AKAP79 (Fig 1K, top row; Fig 1L, gray trace; Supplemental fig 1B). This is consistent with PKAc anchoring to AKAP signaling islands. In contrast, L205R and W196R variants rapidly diffused away from membrane-associated AKAP79, filling the cytosol (Fig 1K, middle and bottom rows; supplemental fig 1B). Quantification of 15 cells from 3 experiments detected significant differences among the variants, including more mobility for PKAc-L205R as compared to PKAc-W196R (Fig 1L & 1M). Similar data were obtained when complementary experiments were performed using the type I PKA-anchoring smAKAP and RIα (Supplemental fig 1C-E). Taken together, these data indicate that Cushing’s syndrome PKAc variants are not compartmentalized by AKAPs.

To gain further insight into the molecular associations and subcellular localization of L205 and W196R PKAc variants in live cells, we conducted proximity biotinylation proteomic experiments (Fig 2A) (Branon et al., 2018; Smith et al., 2018). We generated stable H295R cell lines expressing low levels (~15% endogenous levels to avoid localization artifacts) of WT or mutant PKAc variants tagged with the miniTurbo biotin ligase (Fig 2B, bottom panel; Supplemental fig 2A & 2B). Live-cell labeling for 1 h led to biotinylation of proteins proximal to the tagged catalytic subunit. Distinct patterns of labeled proteins associated with each PKAc-miniTurbo variant were detectable in a neutravidin-HRP western blot (Fig 2B, upper panel; supplemental fig 2A & 2B). Importantly, these studies generated a “proximitome” rather than an “interactome” for each PKAc variant.

**Figure 2.**
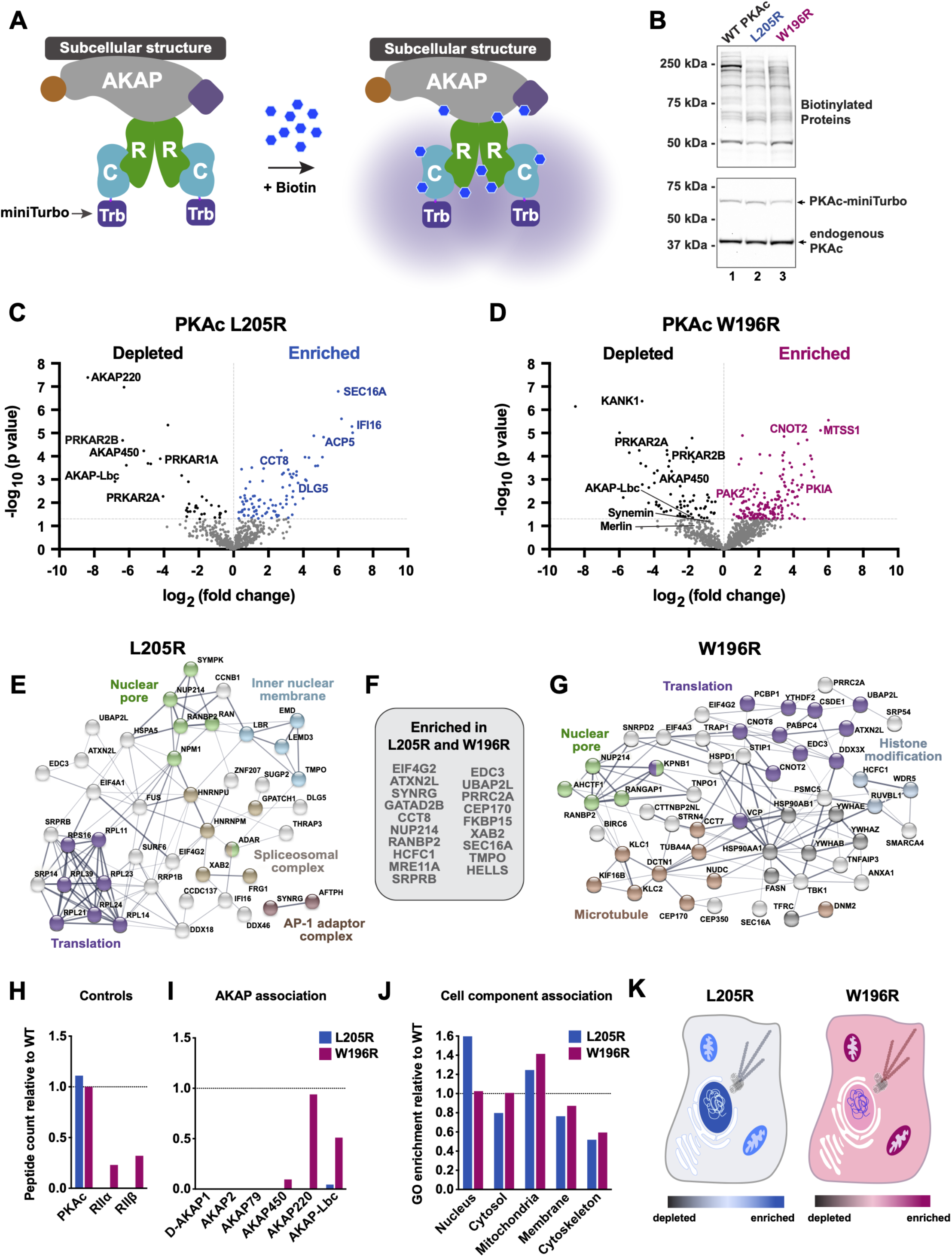
Proximity proteomics identifies distinct associations among PKAc variants. A) Model of PKAc tagged with the biotin ligase miniTurbo and associated with an AKAP signaling island in live cells. Upon application of biotin, proteins in proximity (5-10 nm) to PKAc are biotinylated. B) Immunoblots of lysates from stable H295R cell lines after proximity biotinylation. Neutravidin-HRP signal (top) shows banding differences among the conditions. PKAc signal (bottom) shows comparable expression of tagged kinase at low levels versus endogenous PKAc. Modest expression was used to avoid aberrant localization. C & D) Volcano plots showing distribution of proteins analyzed by mass spectrometry after proximity biotinylation and streptavidin isolation. Blue (L205R) and red (W196R) dots indicate proteins enriched in the mutant condition. Black dots indicate proteins underrepresented as compared to WT. Proteins with a corrected p-value lower that 0.05 are shown in gray. Data from 4 biological replicates. E-G) STRING network depictions of selected enriched proteins in L205R (E) and W196R (G). Between is a list (F) of proteins identified as enriched in both mutant conditions. H) Quantitation of association with PKA holoenzyme components. PKAc represents likely self biotinylation. I) Quantitation of association with AKAP signaling island components as determined by peptides identified relative to the WT condition. J) Gene ontology (GO) enrichment scores for cell component relative to WT. K & L) Graphical depictions of J.

Proteomic analysis identified both shared and distinct pools of proteins between each PKAc Cushing’s variant and the native kinase (Fig 2C and 2D). Importantly, quantitative bioinformatic analyses of each PKAc variant proximitome revealed reduced association with AKAPs and R subunits (Fig 2C and 2D, black dots). Proteins enriched in the mutant conditions (Fig 2C, blue dots; Fig 2D, red dots) were analyzed using STRING to identify network enrichments and gene ontology (GO) cell component annotations (Szklarczyk et al., 2019). Scrutiny of the L205R variant dataset identified enriched subnetworks for cellular processes such as nuclear pore export, protein translation, and spliceosome functions (Fig 2E). Inspection of the PKAc-W196R proximitome also identified components involved in nuclear pore function and translation, as well as distinct enrichment for histone modification and melanosome protein subnetworks (Fig 2G). Comparison between L205R and W196R variant proximitomes revealed nineteen shared, significantly enriched proteins (Figs 2F). Components of the PKA holoenzyme served as positive and negative controls. Accordingly, we detected equivalent self-labeling among PKAc variants and deficient labeling of R subunits (Fig 2H). Loss of association with AKAPs provides further evidence for displacement from AKAP signaling islands in both PKAc-L205R (blue) and PKAc-W196R (red) datasets (Fig 2I). This highlights mechanistic differences between the Cushing’s mutants that are evident in live-cell assays but not immunoprecipitations (Fig 1E & 1F).

By comparing GO enrichment scores for cell component association, we were able to quantify representative differences in the subcellular distribution of PKAc variants (Fig 2J & 2K). PKAc-L205R shows strong enrichment for nuclear components (Fig 2J & 2K, blue). Both mutants exhibit increased proximity to mitochondria components and decreased cytoskeletal associations (Fig 2J & 2K). Taken together, our proximity proteomic results suggest that both Cushing’s PKAc variants exhibit reduced association with AKAPs and appear to be redistributed to different subcellular compartments.

Previous reports have indicated that PKAc-L205R is a tonically active but catalytically inefficient kinase (Lee et al., 2016; Luzi et al., 2018). Moreover, the biochemical measurements and proximity proteomics screens in figures 1 and 2 imply that there may be differences in the mechanisms of action of the PKAc-L205R and PKAc-W196R variants. In vitro activity measurements using purified kinases with Kemptide as a substrate in a real-time kinetic assay (Byrne et al., 2016) confirmed that PKAc-L205R is a less catalytically efficient enzyme than its WT counterpart, requiring 20-fold more kinase to achieve 50% substrate phosphorylation (Fig 3A, blue; supplemental fig 3A). In contrast, substrate phosphorylation curves for PKAc-W196R and the WT kinase were virtually identical (Fig 3A, red and black). ATP incorporation is largely similar among WT and mutants (supplemental fig 3B). Interestingly, mass spectrometry analysis of each PKA variant revealed stark differences in their autophosphorylation status (Fig 3B). Up to 11 autophosphorylation sites were fully occupied on WT PKAc, with W196R showing decreased levels on most sites (Fig 3B). Surprisingly, the PKAc-L205R mutant was almost completely unphosphorylated (Fig 3B). This result provides molecular context to explain the faster migration of PKAc-L205R in gel electrophoresis (supplemental fig 3A). This finding is consistent with evidence that PKAc-L205R is a less stable enzyme, as serine or threonine phosphorylation contributes to the structural integrity of proteins. Accordingly, thermal stability analysis revealed that both Cushing’s mutants are compromised, with unfolding observed at lower temperatures when compared to the WT kinase (Fig 3C). These data highlight physiochemical differences between PKAc-L205R and W196R Cushing’s syndrome variants and illuminate possible explanations for specific mutant kinase behaviors.

**Figure 3.**
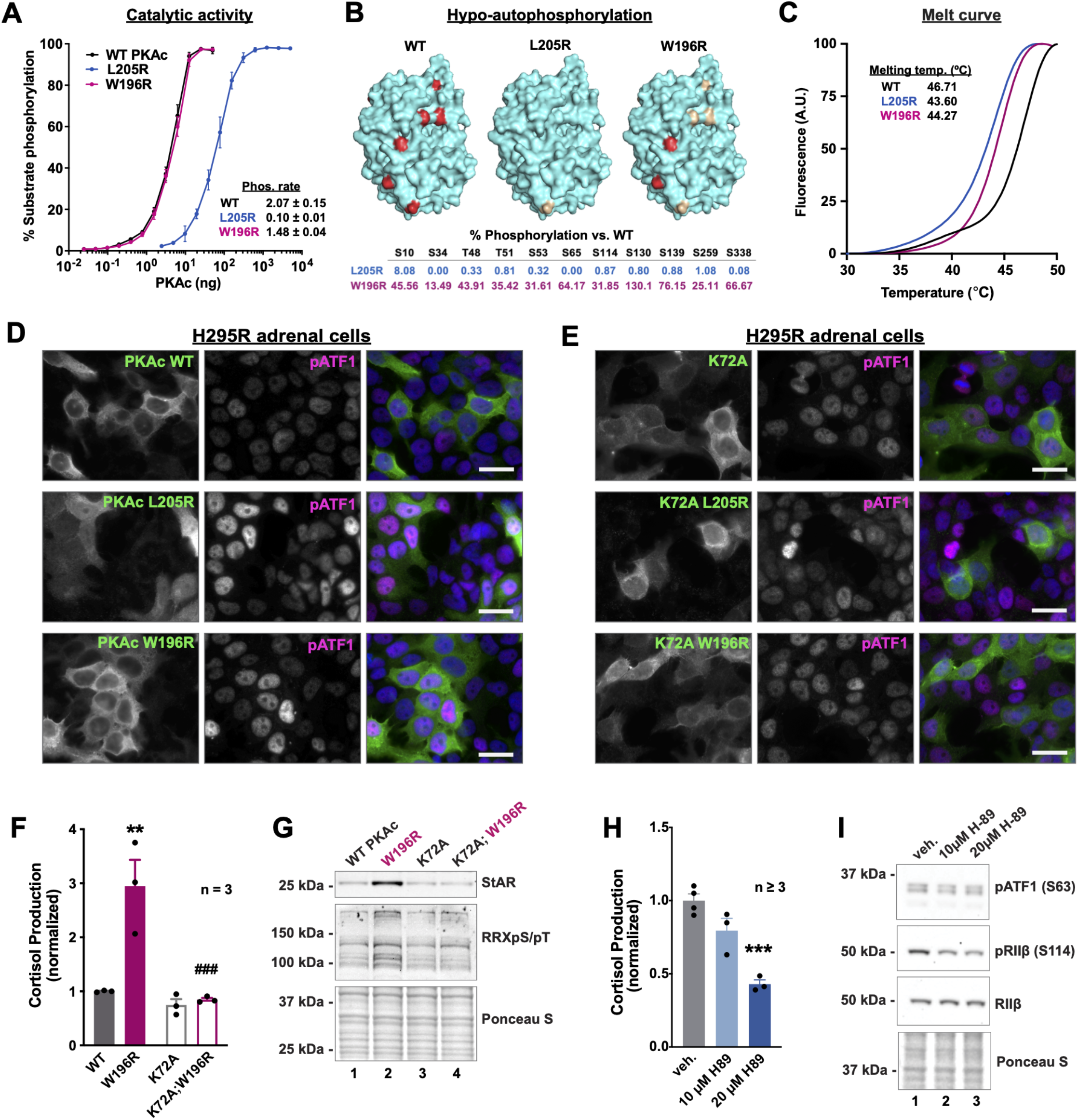
Catalytic activity differs between Cushing’s mutants and is required for elevated stress hormone synthesis. A) Catalytic activity of recombinant WT and mutant PKAc toward a fluorescently labeled peptide substrate. Data is % substrate phosphorylation (mean and SD) after 5 mins assay time. n= 4. Relative rates of activity were determined per ng of enzyme as pmol phosphate incorporation per min. Calculated from assays containing 0.8 ng WT and W196R PKAc and 19.5 ng L205R where linear rates of phosphorylation were detected. B) Structural depictions (top) and quantification (bottom) of relative change in abundance of phosphorylated sites in comparison to WT PKAc. Bright red ≥ 66.67% WT levels. Gold = 1 - 66.66% WT levels. n = 3. C) Normalized thermal melt curves for WT and mutant PKAc variants. D & E) H295R adrenal cells expressing V5-tagged WT or mutant PKAc (D) or kinase dead versions of these carrying the K72A mutation (E) were stained with V5 and pATF1 antibodies. Scale bars = 20 μm. F) Cortisol measurements from H295R cells show that kinase activity is necessary for the mutants effects on cortisol production. **p-value ≤ 0.01 relative to WT and ### p-value ≤ 0.001 relative to W196R, both corrected for multiple comparisons using Sidak method after 1-way ANOVA. n = 3. G) Immunoblot of H295R lysates infected with PKAc variants as listed and probed for steroidogenic acute regulatory protein (StAR) and phosphorylated RRXS/T motif. Representative of three replicate experiments. H) Cortisol measurements from H295R cells after 1 h incubation with vehicle or the PKA inhibitor H89. ***p-value ≤ 0.001. Corrected for multiple comparisons using Sidak method. n ≥ 3. I) Immunoblot of H295R lysates after 1 h incubation with vehicle or H89. Representative of at least 3 experimental replicates.

We next used immunofluorescent techniques to establish the subcellular location of PKAc Cushing’s variants in H295R adrenal cells. While both WT and PKAc-W196R accumulate in the cytosol, the L205R variant displays a more pronounced nuclear localization (Fig 3D, left panels). This finding corresponds with our proximity proteomics data (Fig 2J). In addition, expression of either mutant kinase correlated with enhanced phosphorylation of the canonical PKA substrate ATF1 as detected by phosphopeptide antibodies against the conserved pSer 63 site (Fig 3D, middle panels). Importantly, ATF1 pSer 63 phosphorylation was attenuated when parallel studies were conducted in H295R adrenal cells expressing K72A kinase-dead mutants (Fig 3E). The same effects on substrate phosphorylation were observed for phospho-CREB in ATC7L mouse adrenal cells (supplemental fig 3C & 3D) (Ragazzon et al., 2006). Further characterization of the PKAc-W196R variant involved measuring cortisol release form H295R (Fig 3F). Control experiments confirmed that expression of a kinase dead mutation in W196R rescued overproduction of cortisol back to WT levels (Fig 3F). Biochemical studies in these cells demonstrated that introduction of the kinase dead mutation in PKAc-W196R abolishes increases in PKA substrate phosphorylation (Fig 3G, middle panel, lanes 2 & 4) as well as increases in levels of the mitochondrial cholesterol importer steroidogenic acute regulatory protein (StAR) (Fig 3G, top panel, lanes 2 & 4).

Finally, we tested PKA-dependence of cortisol production in healthy conditions. Inhibition of WT kinase with the PKA inhibitor H89 for 1 h was sufficient to decrease cortisol production (Fig 3H), suggesting that acute PKA-dependent mechanisms contribute to steroid synthesis. Accompanying immunoblot analyses from these cells demonstrate modest reductions in pATF1 levels as well as autophosphorylation of RIIβ (Fig 3I). Replication of H89 experiments in ATC7L cells exhibit similar effects, though indicate a greater sensitivity to the inhibitor (supplemental fig 3E & 3F). Together, these data demonstrate differences in activity and localization between L205R and W196R Cushing’s mutations. Furthermore, kinase-dead experiments confirm that mutant kinase activity is necessary for acute mechanisms of cortisol overproduction in the disease.

To test whether acute activation of WT kinase is sufficient to elevate stress hormone synthesis, we administered cAMP analog drugs known to activate PKA (Fig 4A) (Dostmann et al., 1990; Poppe et al., 2008; Sandberg et al., 1991). Stress hormone measurements demonstrated remarkable differences between the two analog conditions in H295R (Fig 4B) and ATC7L (supplemental fig 4A) cells. At all concentrations tested, the BIMPS/Bnz cAMP drug cocktail failed to elevate stress hormone levels (Fig 4B; green; supplemental fig 4A, cyan). In contrast, application of Sp-cAMPS induced dosage dependent increases in stress hormone production (Fig 4B, purple; supplemental fig 4A, magenta). Control experiments used the adenylyl cyclase activator forskolin (10uM) and the phosphodiesterase inhibitor 3-isobutyl-1-methylxanthine (IBMX; 75uM) to induce supraphysiological second messenger production (Fig 4B & supplemental fig 4A, gray).

**Figure 4.**
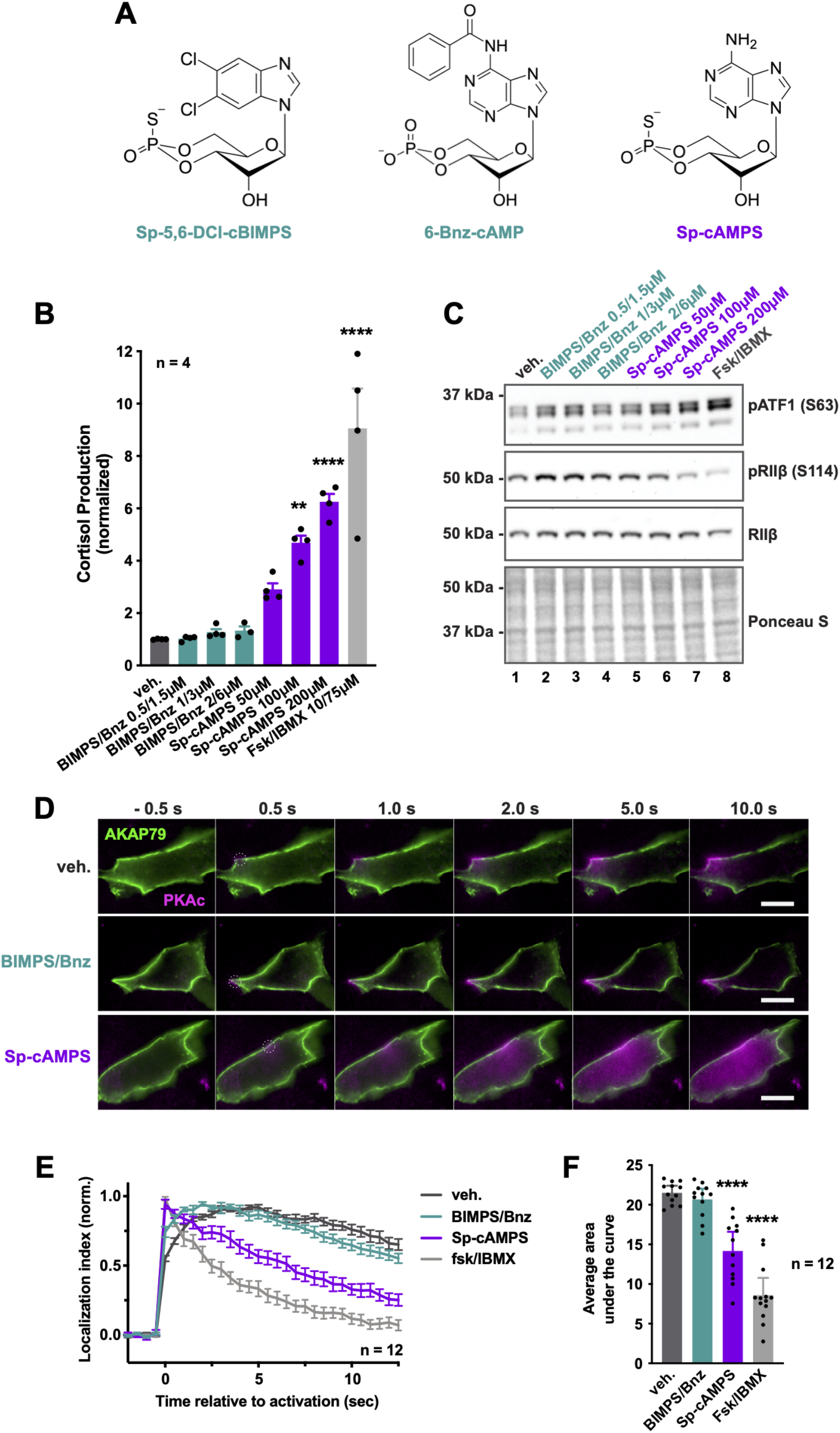
PKA activation alone is not sufficient for stress hormone overproduction. A) structural diagrams of cAMP analogs used in this study. B) Cortisol measurements from H295R cells treated for 1 h with vehicle or increasing concentrations of PKA-activating drugs. ***p ≤ 0.001, ****p ≤ 0.0001, corrected for multiple comparisons with Dunnett method. n = 4. C) Immunoblot of H295R lysates treated for 1 h with vehicle or increasing concentrations of PKA-activating drugs probed for pATF1, pRIIβ, and RIIβ. Representative of 3 experimental replicates. D) Photoactivation microscopy timecourse of H295R cells expressing AKAP79-YFP, RII-iRFP, and WT PKAc tagged with photoactivatable mCherry. Top, vehicle only control; middle, 1 μM Sp-5,6-DCl-cBIMPS in combination with 3 μM 6-Bnz-cAMP; bottom, 100 μM Sp-cAMPS. White circle indicates region of photoactivation. Scale bar = 10 μm. E) Amalgamated data from 3 experimental replicates. n = 12. F) Integration of values from E. ****p ≤ 0.0001, corrected for multiple comparisons with Dunnett method after 1-way ANOVA.

Parallel studies measured ATF1 and RII phosphorylation to test whether the distinct cAMP analog treatments caused differential PKA activation (Fig 4C & supplemental fig 4B). ATF1 phosphorylation appeared equivalent for low and medium concentrations between the two analog conditions tested, suggesting both treatments sufficiently activated the kinase (Fig 4C & supplemental fig 4B, top panels). Immunoblot detection of pSer 114 on RIIβ was used as an index for kinase activation within the context of the anchored PKA holoenzyme (Fig 4C, second panel & supplemental fig 4B, middle panel). Phospho-RIIβ detection was prominent at all concentrations of BIMPS/Bnz (Fig 4C & supplemental fig 4B, lanes 2-4). Conversely, RIIβ ser114 phosphorylation decreased with increasing dosage of Sp-cAMPS, mimicking effects of forskolin/IBMX (Fig 4C & supplemental fig 4B, lanes 5-8). This observation is consistent with the possible release of PKAc from the anchored holoenzyme and its movement away from sites proximal to RIIβ. Based on these endocrine, pharmacological, and biochemical studies, we hypothesized that application of the BIMPS/Bnz cAMP analog cocktail activates PKAc within the context of anchored holoenzymes, whereas delivery of Sp-cAMPS mimics the action of our Cushing’s syndrome mutants by causing the displacement of active PKAc from AKAP signaling islands. This would imply that displacement, as well as activation, is necessary to drive the disease.

To test this postulate, we turned back to our live-cell photoactivation/diffusion assay (Fig 4D). H295R cells were transfected with AKAP79-YFP (green) to sequester PKA at the plasma membrane. Diffusion of photoactivated WT PKAc (magenta) was minimal in vehicle-only and BIMPS/Bnz cAMP analog-treated (1 μM/3 μM) cells over a timecourse of 10 s (Fig 4D, top panels). However, application of Sp-cAMPS (100 μM) promoted release and diffusion of WT PKAc from sites of photoactivation (Fig 4D, bottom panel). As expected, supraphysiological stimulation of cAMP synthesis with forskolin/IBMX induced a complete loss of kinase localization to AKAP signaling islands (supplemental fig 4C). Quantification from multiple experiments confirms the results (Fig 4E and 4F). Together these experiments suggest that activation of PKAc alone is not sufficient to elevate stress hormone production. Rather, these data support a model where displacement of PKAc from AKAP signaling islands and redistribution of the active kinase is necessary to trigger the aberrant cortisol overproduction that is emblematic of Cushing’s syndrome.

On the basis of our findings so far, we postulate that displacement of active PKAc mutants from AKAP signaling islands underlies Cushing’s syndrome (Fig 5A). This hypothesis presupposes that correcting the displacement of Cushing’s PKAc variants should abolish their effects on stress hormone levels. These rescue experiments were achieved by tethering L205R or W196R mutants to RII with a peptide linker (Fig 5B). Constructs where the linker contains a P2A self-cleaving peptide served as controls. An advantage of this latter system is that it ensured equivalent translation of both PKAc and RII subunit protein components (Liu et al., 2017). We established proof of principle for this system in U2OS cells expressing the plasma membrane-localizing scaffold AKAP79 (Fig 5C & 5D). Immunofluorescence detection of AKAP79 (green, left) and PKAc (magenta, middle) demonstrated the subcellular distribution of the anchoring protein and kinase variants respectively. Under control conditions where RII and PKAc are not fused, only native PKAc colocalized with AKAP79 (Fig 5C, top row). In contrast, the free PKAc-L205R and PKAc-W196R mutants filled the cytosol and were displaced from AKAP79 signaling islands (Fig 5C, mid and bottom rows). Importantly, immunofluorescence detection established that each RII-PKAc fusion construct (magenta) co-localized with AKAP79 (green, Fig 5D). Thus, fusion of PKAc-L205R or PKAc-W196R to RII subunits restores the sequestering of PKAc within AKAP signaling islands.

**Figure 5.**
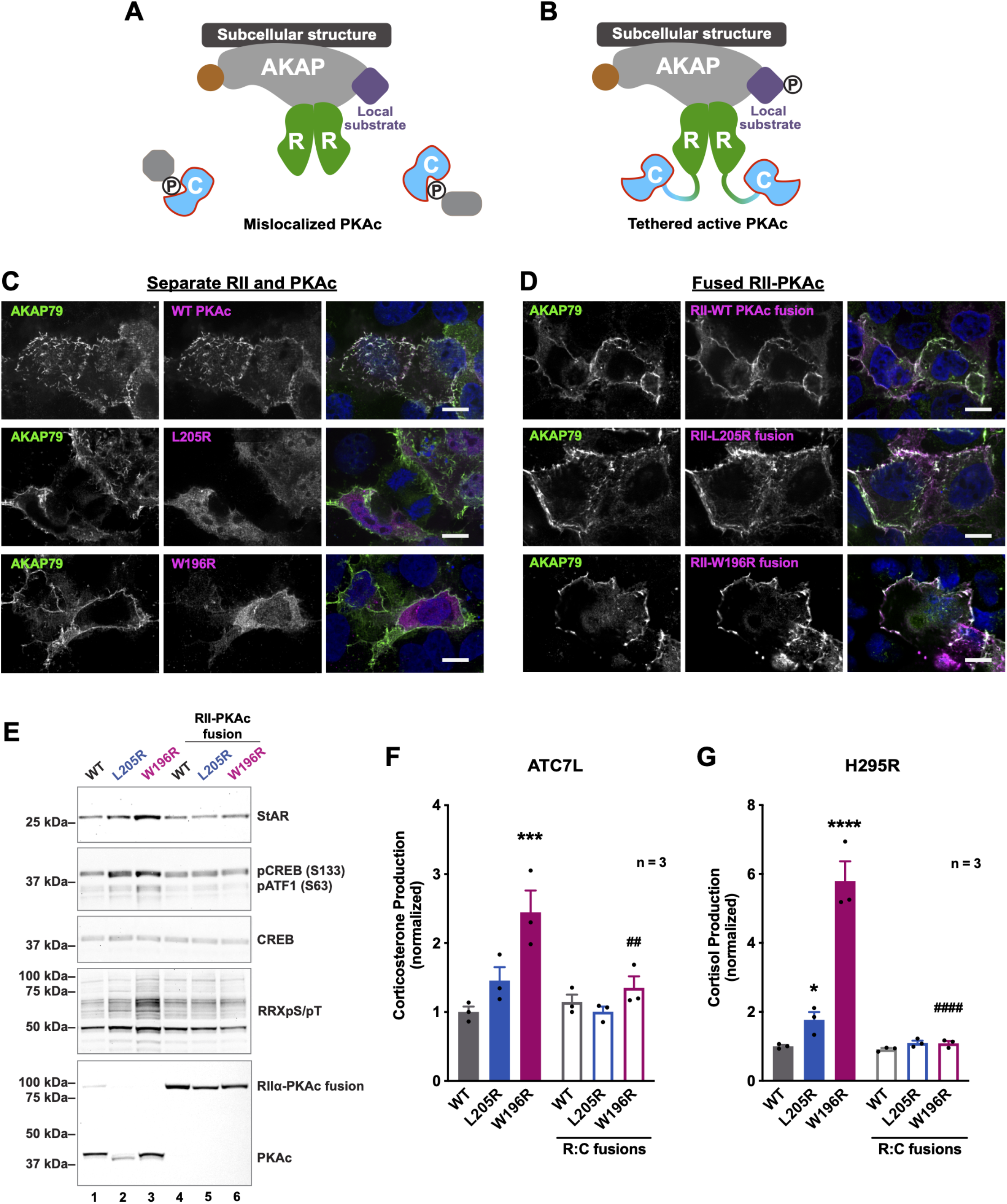
Rescuing AKAP association of Cushing’s mutants restores normal stress hormone production. A) Model of PKAc Cushing’s mutant displaced from AKAPs and catalytically active. B) Model of our strategy to tether these active kinase mutants to AKAP signaling islands by fusing constructs of RII and L205R or W196R. C) U2OS cells expressing AKAP79 (green) along with a self-cleaving construct of RII-P2A-PKAc-V5 (PKAc in magenta). Scale bar = 10 μm. D) Same as C but with fused constructs of RII-PKAc (magenta). Scale bar = 10 μm. E) Immunoblot of ATC7L adrenal cell lysates. Conditions expressing Cushing’s mutants separate from RII (lanes 2 & 3) showed increased signals for StAR, pCREB, pATF1, and phosphorylated RRXS/T. Conditions expressing fused RII-mutant PKAc constructs exhibited the same levels as WT. Representative of 3 replicates. F) Corticosterone measurements from ATC7L adrenal cells expressing either separate or fused RII and PKAc. ***p ≤ 0.001 versus WT, ##p ≤ 0.01 versus W196R unfused condition. Corrected for multiple comparisons using the Sidak method after 1-wau ANOVA. n = 3. G) Cortisol measurements from H295R adrenal cells expressing either separate or fused RII and PKAc. ****p ≤ 0.0001 versus WT, ####p ≤ 0.0001 versus W196R unfused condition. Corrected for multiple comparisons using the Sidak method after 1-wau ANOVA. n = 3.

To test whether localizing mutant kinases to AKAPs in adrenal cells can correct stress hormone overproduction, we expressed control or fusion constructs in ATC7L cells. Overexpression of AKAP79 was omitted from these experiments to ensure that each fusion construct would localize PKA activity to endogenous AKAP signaling islands in these adrenal cells. Immunoblot analyses demonstrated that both unfused Cushing’s mutants elevate levels of StAR as compared to WT PKAc (Fig 5E top panel lanes 1-3). Similarly, phosphopeptide antibodies detected elevation in pCREB, pATF1, and RRXS/T phosphorylation for mutant control conditions (Fig 5E, second and fourth panels, lanes 2 and 3). Fusion of the mutant kinases to RII restored each of these aberrant signaling events back to baseline levels (Fig 5E, lanes 4-6). Finally, fusion of PKAc-L205R or PKAc-W196R to RII rescued overproduction of stress hormone in both mouse ATC7L and human H295R adrenal cells (Fig 5F & 5G). These data strongly suggest that displacement from AKAP/PKA signaling islands is a necessary component of pathology in Cushing’s syndrome.

Our working premise is that displaced Cushing’s PKAc variants bring about pathologic conditions via altered phosphorylation of substrate proteins within their vicinity. To further elucidate signaling mechanisms dysregulated in the disease, we conducted phosphoproteomics on proximity-biotinylated proteins isolated from H295R cells expressing L205R, W196R and WT versions of PKAc-miniTurbo. Mass spectrometry analysis and volcano plot visualization show phosphopeptides enriched and depleted for each mutant (Fig 6A & 6B). Comparison of the PKAc-L205R phospho-proximitome to WT revealed 48 hits that were significantly depleted, including phosphopeptides for AKAP-Lbc and AKAP220 (Fig 6A, black dots & 6D, left). Likewise, 30 phosphopeptides were found significantly depleted upon scrutiny of the PKAc-W196R phospho-proximitome, also including AKAP-Lbc and AKAP220 (Fig 6B, black dots & 6D, right). The reduced phosphorylation of AKAPs is once again consistent with our biochemical and live-cell imaging evidence that Cushing’s syndrome PKAc variants are displaced from AKAP signaling islands (Figs 1E & K). Eighty phosphopeptides were enriched in the PKAc-L205R phospho-proximitome (Fig 6A, blue dots & 6C, left) and 45 phosphopeptides were enriched in the PKAc-W196R condition (Fig 6B, red dots & 6C, right). Notably, overlap analysis revealed six proteins significantly enriched and seven significantly depleted between mutant datasets (Fig 6C; Supplemental fig 5A). Among the common enriched proteins was insulin receptor substrate 2 (IRS2) (Fig 6C, middle). Because both IRS2 and AKAP-Lbc regulate MAP kinase signaling (De Meyts, 2000; Smith et al., 2010), we hypothesized that perturbed PKAc location in Cushing’s syndrome could dysregulate downstream ERK signaling and lead to increased proliferation and/or cortisol production from adenomas. Further motivation to pursue this postulate came from NetworKIN analysis of phosphosites upregulated in each mutant condition (Fig 6D & 6E) (Horn et al., 2014). Each mutant was found associated predominately with substrates of proline-directed kinases, including cyclin-dependent kinases and the MAP kinase family.

**Figure 6.**
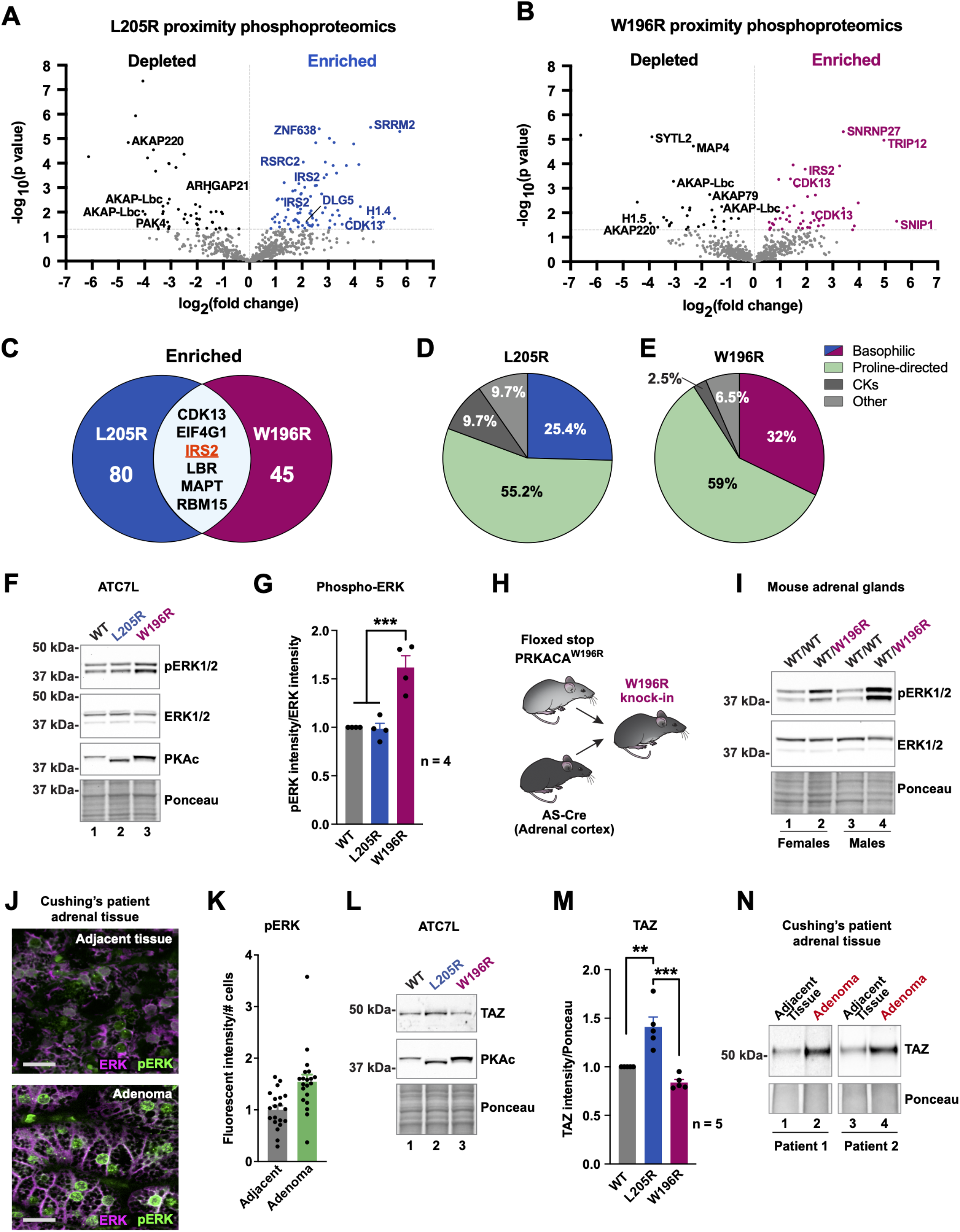
Proximity phosphoproteomics reveal disruption of ERK and Hippo signaling. A & B) Proximity phosphoproteomics of PKAc-miniTurbo biotinylated H295R samples from L205R (A) and W196R (B) vs WT conditions. Phosphopeptides depleted in the mutant conditions are shown as black dots. Enriched phophopeptides are shown in blue (L205R) and red (W196R). C) Venn diagram showing significantly enriched phosphopeptides identified in both mutant conditions. D & E) NetworKIN kinase prediction for phosphosites enriched in L205R (D) and W196R (E) conditions. F) Immunoblot of ATC7L cells expressing PKAc variants. Increased ERK phosphorylation is evident in the W196R condition. G) Quantitation of F. ***p≤0.001. Analyzed with one-way ANOVA and Sidak multiple comparisons correction. n = 4. H) Schematic depicting cross of a conditional PRKACA^W196R^ knock-in mouse with an adrenal cortex-specific AS-Cre line, yielding a heterozygous adrenal cortex-specific W196R mutant mouse. I) Immunoblot of adrenal gland lysates from female (lanes 1 & 2) and male (lanes 3 & 4) littermate mice. Mutant mice show increased phospho-ERK signal compared to littermate controls. J) Immunostaining for ERK (magenta) and phospho-ERK (green) of resected tissue from adenoma (bottom) and adjacent adrenal gland (top) of a Cushing’s syndrome patient. scale bar = 20 μm. K) Quantitation of fluorescent signal for phospho-ERK. 10 images each per condition from 2 patients. L) Immunoblot of ATC7L cells expressing PKAc variants. TAZ levels are elevated in PKAc-L205R but not PKAc-W196R. M) Quantitation of K. ***p≤0.001, **p≤0.01. Analyzed by one-way ANOVA with multiple comparisons corrected by Sidak method. n = 5. N) Immunoblot of resected Cushing’s syndrome tissue (paraffin-embedded sections) from 2 patients. TAZ is elevated in adenoma versus adjacent tissue.

To test whether mutant PKAc expression impacted ERK pathways, we initially probed lysates from ATC7L adrenal cells expressing WT or mutant PKAc. Immunoblot detection of phospho-ERK served as an index of mitogenic kinase activity. Although total ERK levels appeared unchanged from WT in mutant conditions (Fig 6F, second panel), phospho-ERK was elevated in the W196R condition (Fig 6F, top panel lane 3 & 6G). Surprisingly, phospho-ERK levels were unaffected in the presence of PKAc-L205R (Fig 6F, top panel lane 2 & 6G). Next, we tested whether embedding Cushing’s mutants within AKAP signaling islands could rescue ERK phosphorylation. Upon expression of the fused RII-PKAc-W196R kinase, phospho-ERK levels were returned to normal levels (Supplemental fig 5B, top panel lane 6). Fusion of native PKAc or L205R had no effect on immunoblot detection of phospho-ERK (Supplemental fig 5B, lanes 4 & 5). We next wanted to test this in a mouse model of heterozygous W196R expression in the adrenal cortex. Mice containing a conditional allele of PKAc-W196R were crossed to an adrenal cortex-specific Cre line to yield mice heterozygous for the Cushing’s mutation (Fig 6H; Supplemental fig 5C) (Freedman et al., 2013; Niswender et al., 2005). Testing adrenal lysates from both male and female sibling pairs showed increased ERK activation in adrenal glands of mutant mice (Fig 6I, top panel). Total ERK levels were unchanged between WT and mutant mice (Fig 6I, second panel). Paraffin embedded sections of adrenal adenoma and adjacent tissue from Cushing’s patients were stained for phosphorylated (green) and total (magenta) ERK (Fig 6J). Analyses of ten images from each patient revealed a >50% increase of phosoho-ERK signal in adenoma versus adjacent tissue (fold change 1.55 ± .14 SEM) (Fig 6K). These results indicate that ERK activation is enhanced by mutant PKAc variants found in Cushing’s syndrome.

As previously stated, PKAc-L205R phospho-ERK levels were unchanged in adrenal cells (Fig 6G). Upon further scrutiny of our proteomic data, we identified discs large homolog 5 (DLG5) as specifically enriched in the presence of PKAc-L205R (Fig 2C, blue & 6A, blue). Three phosphopeptides were increased in the mutant samples, including the consensus PKA site (R-R-L-S) at residues 1663-1666. DLG5 controls the Hippo pathway, which regulates YAP/TAZ transcription factors and is an evolutionarily conserved signaling mechanism for cell survival and proliferation (Kwan et al., 2016). Furthermore, dysregulated Hippo signaling has been observed in multiple forms of cancer (Yu and Guan, 2013). Upon measuring levels of YAP/TAZ in ATC7L cells, we found upregulated levels of TAZ in PKAc-L205R cells (Fig 6L, lane 2 & 6M, blue). This effect was not observed in cells expressing PKAc-W196R (Fig 6L, lane 3 & 6M, red). Furthermore, immunoblots of Cushing’s syndrome patient samples showed an upregulation of TAZ in adenoma versus adjacent tissue (Fig 6N). Remarkably, these studies demonstrate that these similar Cushing’s syndrome mutants potentiate distinct mitogenic pathways.

## Discussion

We have discovered three fascinating new details about the molecular pathology of Adrenal Cushing’s syndrome. The PKAc-L205R and W196R variants display different physiochemical properties, they are differentially compartmentalized, and each mutant kinase promotes cortisol overproduction via a distinct signaling mechanism. At first glance, the PKAc-L205R and PKAc-W196R variants appear virtually identical. Both disease-causing mutations occur within the same exon of the *PRKACA* gene and promote amino acid changes only nine residues apart. However, protein melting curves in figure 3C indicate that both Cushing’s kinases are more susceptible to protein unfolding at lower temperatures than WT PKAc. Thermal instability of PKAc-L205R or PKAc-W196R could play a part in the etiology of Cushing’s, since factors that influence protein folding and degradation have been implicated in other PKA pathologies (Rinaldi et al., 2019; Turnham et al., 2019). Likewise, auto- and transphosphorylation of the catalytic core contributes to the stability, maturation, and activity of AGC kinases (Baffi et al., 2021; Newton, 2003). The V600E phosphomimetic mutation in BRAF, for example, a common oncogenic driver in melanoma, dramatically elevates basal kinase activity (Davies et al., 2002; Holderfield et al., 2014). Also, multisite phosphorylation of PKC’s is necessary for fully mature kinases, although cancer-associated mutations generally corollate with loss-of-function (Antal et al., 2015; Van et al., 2021). As indicated in figure 1E, the PKAc-L205R has a faster electrophoretic mobility than its counterparts, which is a classic hallmark of reduced phosphate incorporation. Quantitative mass spectroscopy data in figure 3B provides compelling proof of this phenomenon. While the native kinase is fully phosphorylated on eleven sites and PKAc-W196R exhibits sub-stoichiometric incorporation of phosphate at most sites, the L205R variant is almost totally devoid of phosphate. Interestingly, the PKAc-L205R displays a twenty-fold lower rate of catalysis toward in vitro substrates (Fig 3A). This could explain why PKAc-L205R induces lower levels of cortisol production than the PKAc-W196R form. Thus, subtle changes in physiochemical properties of the kinase catalytic core can be amplified by environmental factors into more profound effects on protein kinase A physiology.

A traditional view of cAMP signaling proposes that this diffusible second messenger unilaterally activates PKA throughout the cell. Yet, contrary to this dogma, elegant fluorescent spectroscopy studies show that under physiological parameters cAMP accumulation occurs within nanometer-sized domains (Bock et al., 2021; Musheshe et al., 2018). The existence of cAMP nanodomains argues that the local actions of PKA, EPAC guanine-nucleotide-exchange factors, and Popeye domain-containing proteins are more intricately organized than was originally considered (Amunjela et al., 2019; Beltejar et al., 2017; Laudette et al., 2018; Viña et al., 2021; Yarwood, 2020). This also explains how the ubiquitous second messenger cAMP can simultaneously propagate distinct signaling events within the same cell. This cAMP nanodomain hypothesis is consistent with our own molecular evidence that anchored, intact, and active PKA holoenzymes operate within a 200-400 Å radius of action (Smith et al., 2017; Smith et al., 2013). We call these autonomous cAMP signaling units AKAP signaling islands (Omar and Scott, 2020).

A key discovery of this report is that cortisol overproduction is a consequence of the exclusion of mutant kinases from AKAP signaling islands. This possibility was suggested to us by three lines of evidence. First, live-cell imaging data in figure 1K shows that the PKAc-L205R and W196R variants have a greater intracellular mobility than the WT kinase. Second, quantitative proteomic screening of the PKAc-L205R and PKAc-W196R proximitomes in figure 2 reveal that both Cushing’s variants have reduced interaction with AKAPs. Third, pharmacological studies with cAMP analogs in figure 4 demonstrate that supraphysiological release of native PKAc from AKAPs, but not activation alone, enhances stress hormone production. Taken together, these results converge on the notion that exclusion of PKAc from AKAP signaling islands contributes to hypersecretion of cortisol. This new model is substantiated by rescue experiments in figure 5, wherein sequestration of Cushing’s variants within AKAP signaling islands restores stress hormone production back to baseline levels. Hence, in Cushing’s syndrome, where PKAc acts is more important than when or how the kinase is turned on.

The aberrant phosphorylation events that ensue upon mislocalization of active Cushing’s kinases can have both acute and chronic effects on adrenal cell pathophysiology. Interestingly, cAMP analog experiments in figure 4B imply that pharmacological activation and displacement of endogenous PKAc can drive stress hormone overproduction within an hour. This implies that a sizeable component of mutant PKAc’s pathological impact proceeds via non-transcriptional mechanisms. Nonetheless, pathological cAMP responsive events must also contribute to the chronic tumor growth emblematic of Cushing’s syndrome. Consistent with this notion, data presented in figures 3G and 5E show that the mitochondrial cholesterol transporter steroidogenic acute regulatory protein (StAR) is markedly upregulated by Cushing’s PKAc variants. Remarkably, biochemical and cell-based experiments in figure 6 reveal that the L205R and W196R Cushing’s variants engage different mitogenic signaling pathways. Proximity phosphoproteomics and cell-based validation in figure 6A-G identify upregulated MAP kinase activity in adrenal cells expressing PKAc-W196R. A more stringent test of this hypothesis was to generate an adrenal selective knock-in of PKAc-W196R in mice. This led us to discover enhanced ERK activation in adrenal glands from animals expressing a single allele of this Cushing’s PKAc mutation. In addition, staining of tissue sections from Cushing’s patients in figure 6J detects elevated ERK signaling in adenomas. Importantly, ERK activation was restored to normal levels when the W196R variant was tethered to endogenous adrenal cell AKAPs. This suggests that stress hormone synthesis and ERK activation are co-regulated by Cushing’s PKAc mutants. This is the first report of augmented ERK activation by mutant PKAc in Cushing’s syndrome. Indirect support for this finding comes from recent evidence that the MEK inhibitor PD184352 reduces basal cortisol production from H295R cells (Pereira et al., 2019). Furthermore, AKAP-Lbc, one of the most prevalent AKAPs identified in our proteomic experiments, is a known regulator of MAP kinase signaling. This tallies with recent whole exome sequencing of a Cushing’s syndrome patient that identified an allele of AKAP-Lbc fused to the phosphodiesterase PDE8A in a (Di Dalmazi et al., 2020). At the protein level, this chimeric protein could efficiently terminate local PKA signaling by metabolizing cAMP within AKAP-Lbc signaling islands. Understanding how individual AKAP signaling islands regulate cortisol production and how the ERK pathway overrides this process represent future avenues for therapeutic intervention in Cushing’s syndrome.

Unexpectedly, signaling events that emanate from PKAc-L205R are markedly different. As indicated in figures 6F-N this Cushing’s mutant does not potentiate MAP kinases but instead regulates the Hippo pathway, an evolutionarily conserved signaling cascade that controls organ size and development (Yu and Guan, 2013). A first indication of this remarkable finding was enhanced detection of disks large 5 (DLG5) in the PKAc-L205R proximitome (Figure 2C). This scaffolding protein potently inhibits Hippo signaling (Kwan et al., 2016). Additionally, phosphoproteomic analysis in figure 6A revealed increased phosphorylation of DLG5 in the L205R condition. This led to the discovery in figure 6L that adrenal cells expressing the PKAc-L205R variant contain increased levels of TAZ, a well-studied transcriptional activator of proliferation (Moroishi et al., 2015). Similar effects were observed upon biochemical analyses of resected tissue from Cushing’s adenomas (Fig 6N). Remarkably, TAZ levels were unaffected in cells or tissues expressing PKAcW196R. Thus, Cushing’s syndrome-causing mutations in PKAc that occur nine residues apart mobilize mutually exclusive downstream signaling events. Interestingly, a recent clinical study also reported an inverse correlation between ERK1 and YAP1 levels in breast cancer patients as well as increased YAP1 expression upon ERK silencing (Yu et al., 2019). In conclusion, we have discovered a remarkable molecular mechanism whereby PKAc-L205R and PKAc-W196R variants, that are refractory to AKAP targeting, drive adrenal Cushing’s syndrome via the mobilization of distinct signaling cascades.

## Acknowledgments

The authors would like to thank K. Rosenthal, L. Gabrovsek, J. Nelson, K. Suso, and H. Dahlin for technical assistance and helpful discussions. This work was supported by NIH grants F32DK121415 (M.H.O.), DK119186 (J.D.S.), and DK119192 (J.D.S.), a research grant from the Fibrolamellar Foundation (J.D.S.), funding from BBSRC BB/T018127/1 and BB/S018514/1 (C.E.E. & P.A.E.), and Cancer Research UK C1443/A22095 (C.E.E.). The authors declare no conflicts of interest.

## Supplemental figure legends

**Supplemental figure 1.**
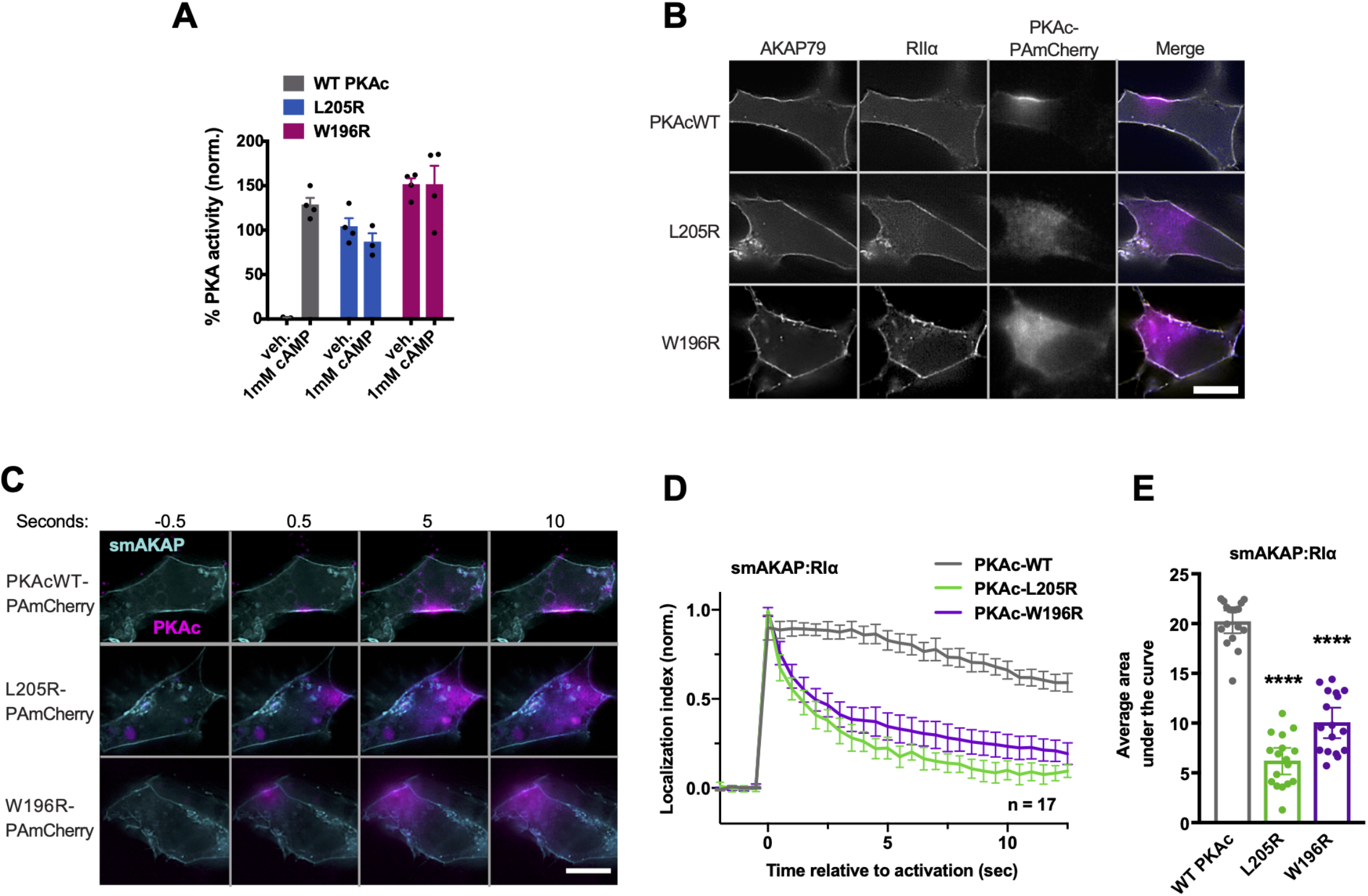
A) Activity of PKAc variants in the presence of RII +/- cAMP. n = 4. B) 3-channel representative images of photoactivation cells demonstrating colocalization of AKAP79 and RII signals. scale bar = 10 μm. C) Photoactivation timecourses in H295R cells with type I PKA. smAKAP-GFP, RI-iRFP, and PKAc tagged with photoactivatable mCherry were expressed. Scale bar = 10 μm. D) Amalgamated quantitation of C. Over time, the majority of WT PKAc (gray) remains localized to smAKAP while L205R (green) and W196R (purple) diffuse away. Localization index = fluorescence intensity at the site of activation divided by intensity at off target sites. Data represents 3 experimental replicates. E Integration of values in D. ****p ≤ 0.0001, corrected for multiple comparisons with Dunnett method after 1-way ANOVA.

**Supplemental figure 2.**
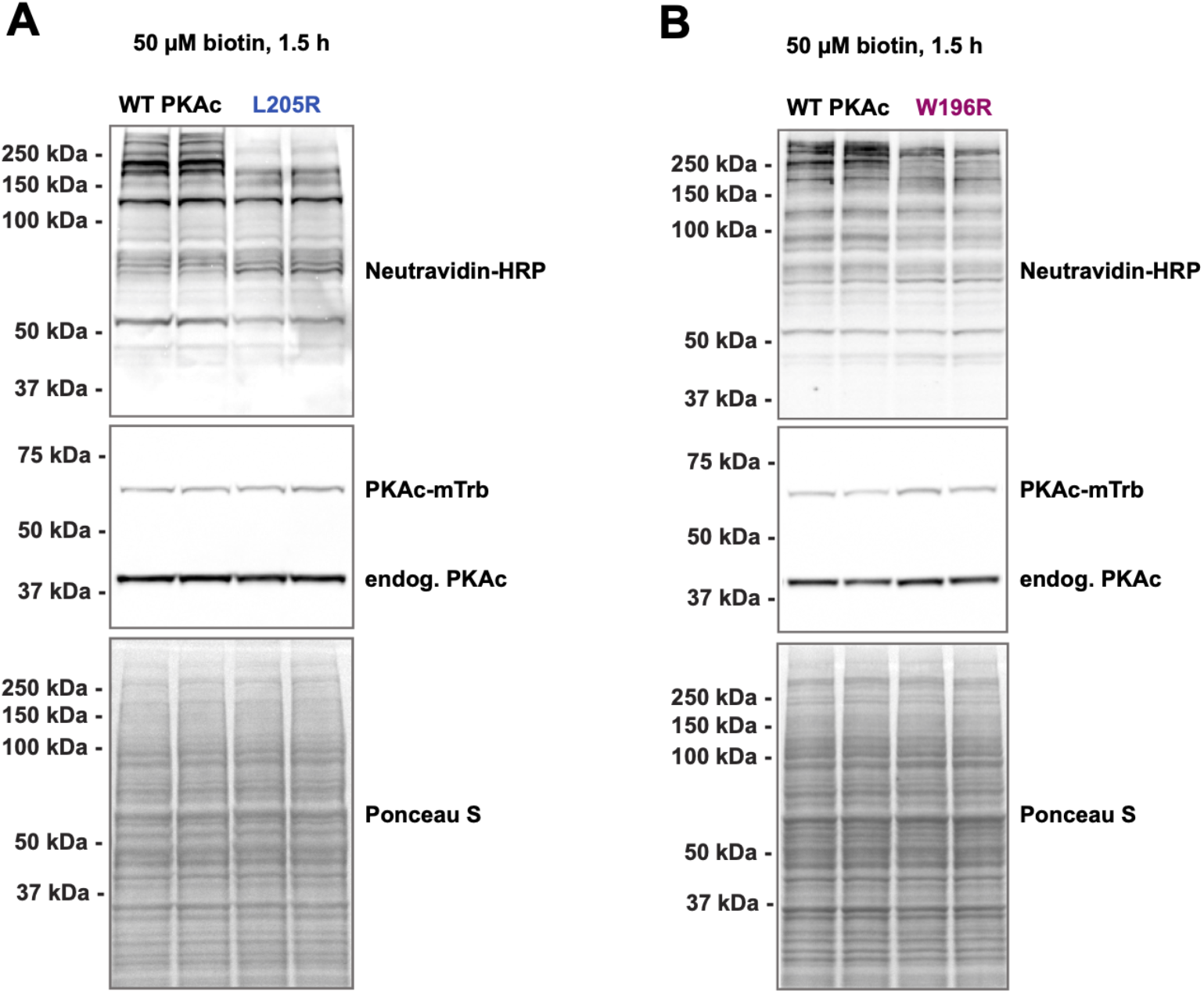
Representative western blots for L205R (A) and W196R (B) variants of PKAc-miniTurbo expressed in H295R cells and incubated with 50 μM biotin for 1.5 h.

**Supplemental figure 3.**
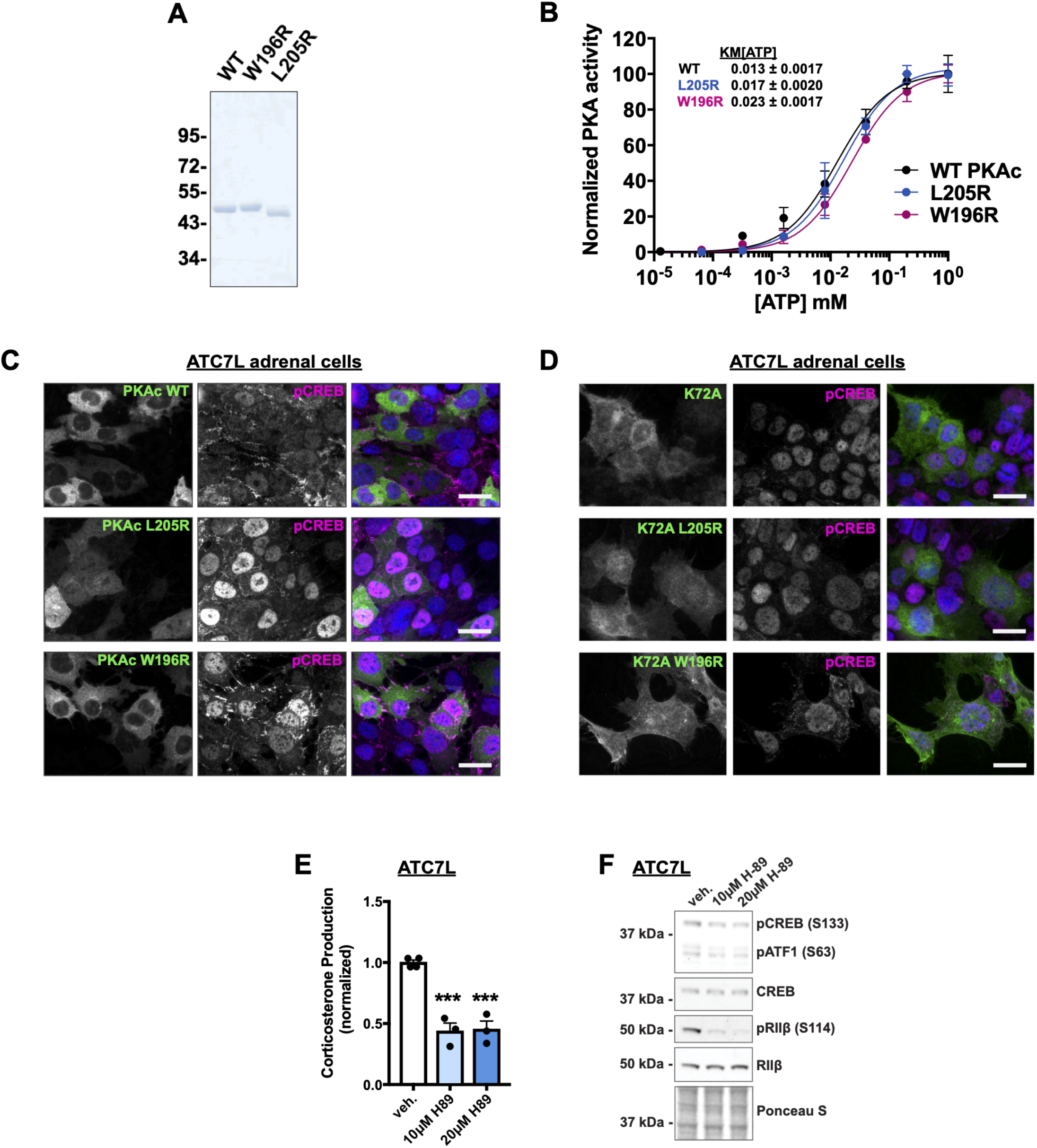
A) SDS-PAGE and coomassie blue staining of purified recombinant WT and mutant PKAc variants. B) KM[ATP] determination for recombinant WT and mutant PKAc. Data shown is PKA-catalyzed peptide phosphorylation (pmol phosphate/min/ng protein) with increasing concentrations of ATP. Normalized to the maximum rate of phosphorylation for each protein. KM[ATP] values (± SE) were calculated from four independent experiments. C & D) ATC7L mouse adrenal cells expressing V5-tagged WT or mutant PKAc (C) or kinase dead (K72A) versions of these (D) were stained with V5 and pCREB antibodies. Scale bars = 20 μm. E) Corticosterone measurements from ATC7L cells after 1 h incubation with vehicle or the PKA inhibitor H89. ***p-value ≤ 0.001. Corrected for multiple comparisons using Sidak method. n ≥ 3. I) Immunoblot of ATC7L lysates after 1 h incubation with vehicle or H89. Representative of at least 3 experimental replicates.

**Supplemental figure 4.**
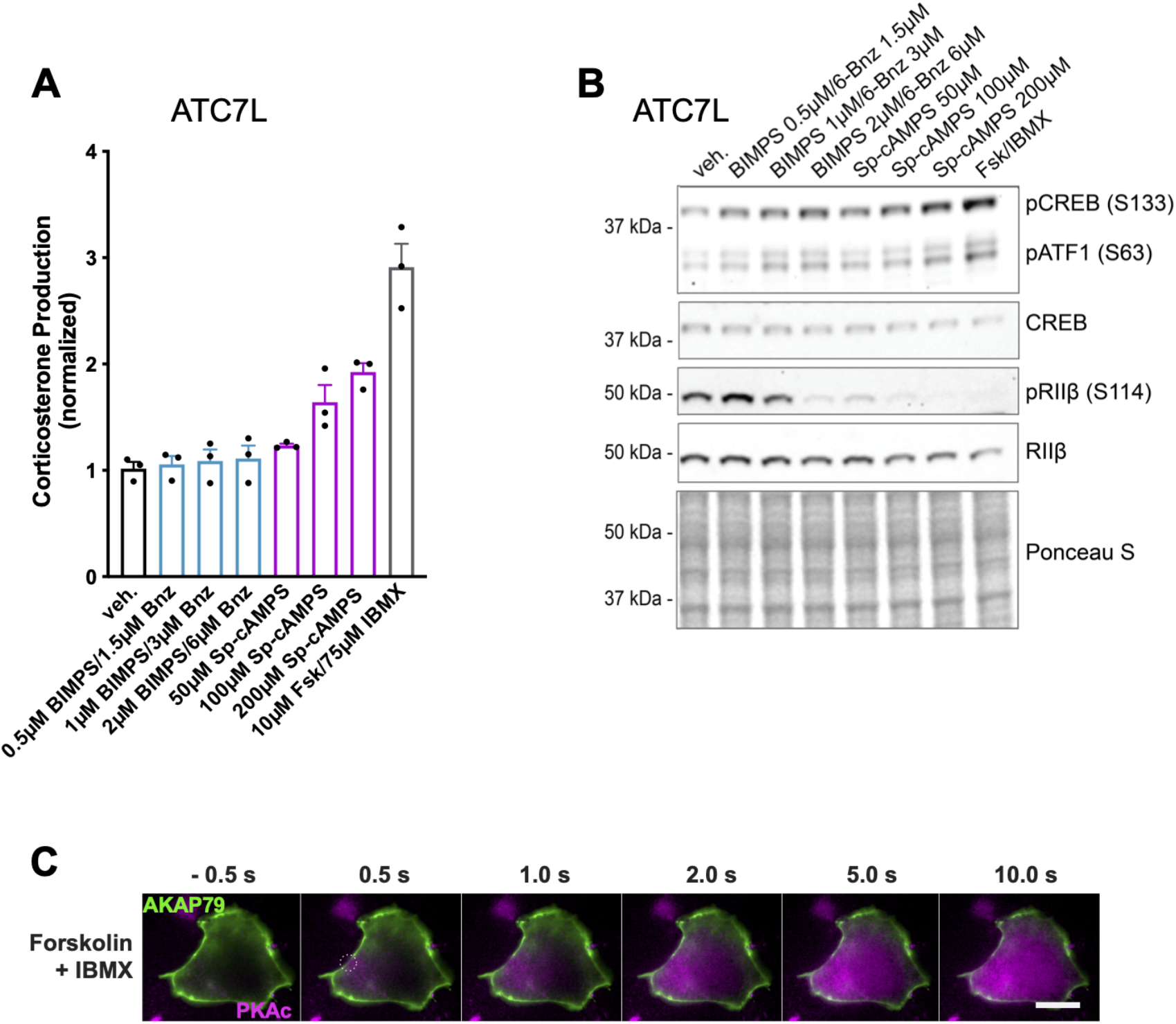
A) Corticosterone measurements from ATC7L cells treated for 1 h with vehicle or increasing concentrations of PKA-activating drugs. n = 3. B) Immunoblot of ATC7L lysates treated for 1 h with vehicle or increasing concentrations of PKA-activating drugs probed for pCREB/pATF1, total CREB, pRIIβ, and RIIβ. Representative of 3 experimental replicates. C) Photoactivation microscopy timecourse of H295R cell expressing AKAP79-YFP, RII-iRFP, and WT PKAc tagged with photoactivatable mCherry in the presence of 10 μM forskolin and 75 μM IBMX. White circle indicates region of photoactivation. Scale bar = 10 μm.

**Supplemental figure 5.**
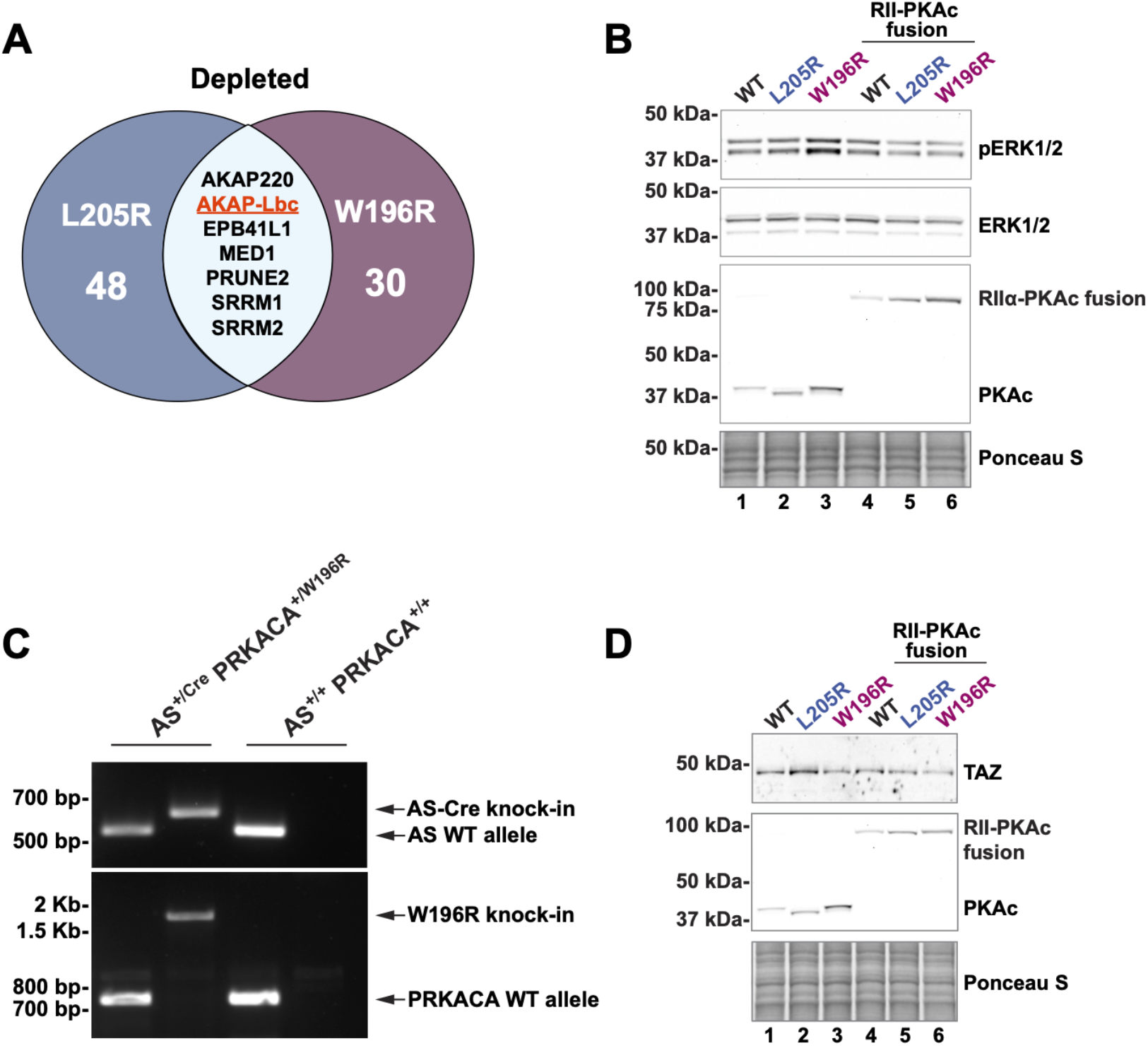
A) Venn diagram of significantly depleted phophopeptides in both mutant conditions. B) Immunoblot of ATC7L cells lysates. Conditions expressing separate RII and PKAc proteins demonstrated increased ERK phosphorylation in the W196R condition. This effect was rescued by fusion of RII and PKAc. C) Agarose gel of genotyping PCR reactions using tail DNA from littermate mutant (left 2 lanes) and WT (right 2 lanes) male mice. D) Immunoblot of ATC7L cells expressing separate (lanes 1-3) or fused (lanes 4-6) RII and PKAc variants.

## Materials and methods

### Protein structure models

Models of PKA catalytic were made using PYMOL. Database files for human (number) and mouse (number) versions of the kinase were used.

### Antibodies

The following antibodies were used in our studies: PKAc BD 610981 (IF, WB); PKA RIIα BD 612243 (WB); GFP Rockland 600-101-215 (IF, IP, WB); AKAP79 custom V089 (WB); PKA RIβ Santa Cruz sc-907 (WB); V5-tag Thermo Fisher R96025 (IF, IP); phospho-CREB/phospho-ATF1 CST 9198 (IF, WB); CREB CST 9104 (IF, WB); Phospho-PKA Substrate CST 9624 (WB); NeutrAvidin-HRP Pierce 31030 (WB); StAR CST 8449 (WB); phospho-RIIβ Santa Cruz sc-136460 (WB); RIIβ BD 610626; phospho-ERK1/2 CST 9101 (IF, WB); ERK1/2 CST 9102 (WB); pan ERK BD 610123 (IF); YAP/TAZ CST 8418.

### Immunoprecipitations

Cell lysates were made using lysis buffer containing 1% TritonX100, 130 mM NaCl, 20 mM NaF, 2 mM EDTA, and 50 mM Tris pH 7.5 (at 4°C) along with 1 mM AEBSF, 10 μM leupeptin, and 1 mM benzamidine. Protein concentrations were adjusted to 1 mg/mL using lysis buffer and precleared by rotating with 20 μL protein G agarose for 30 min at 4°C. Supernatants were then incubated with 1-2 μg of the appropriate antibody overnight. In the morning 30 μL of protein G agarose was added and samples were returned to rotation for 1 h. Samples were washed 3x with lysis buffer and centrifugation at 5000xg, and then dried with a 27G needle before resuspending in 1x PAGE sample buffer. Figures are representative for at least 3 experimental replicates.

### Immunoblotting

Cell lysates were made using RIPA lysis buffer (1% NP-40 Tergitol, 0.5% deoxycholate, 0.1% SDS, 130 mM NaCl, 20 mM NaF, 2 mM EDTA, and 20 mM Tris pH 7.5 (at 4°C) along with 1 mM AEBSF, 10 μM leupeptin, and 1 mM benzamidine). For experiments to detect S/T phosphoproteins, 10 mM beta glycerophosphate was added. Concentrations were measured using BCA. Gels were loaded with 10-30 μg protein after heating for 10 min at 80°C. Proteins were transferred to nitrocellulose or PDVF, incubated with ponceau S to measure total protein loading, blocked in 5% milk TBST for 30 min at RT, and probed with antibodies in 5% BSA TBST overnight at 4°C. Membranes were then incubated with secondary antibodies conjugated to HRP diluted in 5% milk TBST for 1 h at RT. Signals were visualized with chemiluminescence on an Invitrogen iBright Imaging System.

### Stress hormone measurements

Cells were washed once in PBS and media was changed 1.5 h before harvest. Harvested media samples were snap frozen until assayed. To assay, samples were diluted 5 to 10-fold and subjected to measurement using Enzo Cortisol or Corticosterone ELISA kit.

### Photoactivation microscopy

NCI-H295R or HEK293T cells were grown in glass bottom dishes and transfected using Lipofectamine 3000 24-48 h before imaging. Plasmids encoding AKAP79-YFP, D-AKAP1-GFP, smAKAP-GFP, RIIα-iRFP, RIα-iRFP, and either WT, L205R, or W196R tagged with photoactivatable mCherry were used. Imaging was performed using a GE OMX SR system. Exposure and laser intensity were optimized for each experimental replicate and held constant among experimental conditions. Photoactivation laser duration was kept under 50 milliseconds to activate a discrete area with minimal spread in the first image collected after activation. Images were collected at 2 Hz in 3 channels. A baseline of 4 images was taken prior to activation of the PKAc fluorophore. Cells were selected for imaging only when R-iRFP signal was clearly colocalized with the AKAP. Secondary screening for this was performed posthoc as well. Timecourses were measured using ImageJ analysis software (FIJI). A localization index (intensity of the activated region divided by intensity of cytosolic region at least 6 μm distal) was used to interrogate change in fluorescent signal localization over time (mobility).

### Proximity biotinylation and sample preparation for MS

Stable NCI-H295R adrenal cell lines were made using lentivirus encoding a tetracycline-responsive promoter and variants of PKAc tagged with V5 and miniTurbo biotin ligase at the c-terminus. Induction with doxycycline was optimized to yield a subtle overexpression of the bait constructs at 15% of endogenous PKAc levels, as determined by PKAc immunoblotting and quantification using Image J. miniTurbo-tagged variant expression was induced for 48 h before application of 50 μM biotin in DMSO. Cells were incubated for 1 h at 37°C, washed 2x for 1 min using 10 mL PBS to deplete excess biotin, and then lysed using RIPA buffer (as described above). Protein concentrations were measured by BCA and samples were diluted to 1 mL of 0.5 mg/mL in RIPA and placed in low protein-binding tubes containing 25 μL of Nanolink magnetic streptavidin beads. Tubes were rotated 1 h at RT and placed on a magnet. Supernatant was saved for diagnostics and samples were washed in RIPA 2x, 2 M urea in 20 mM Tris 2x, and 25 mM Tris 2x. For normal mass spec analysis, samples were resuspended in 8 M urea in 100 mM Tris pH 8.5 with 5 mM tris(2-carboxyethyl)phosphine hydrochloride (TCEP) and 10 mM chloroacetamide (CAM) and then incubated at 37°C for 1 h. For phosphopeptide mass spec analysis, samples were resuspended in 20% trifluoroethanol 25 mM Tris pH 7.8 with 5 mM TCEP and 10 mM CAM and incubated at 95°C for 5 min. For digestion, samples were diluted 2-fold with 100 mM TEAB and 1 μg LysC was added before incubation for 2 h shaking at 37°C. Samples were again diluted with 100 mM TEAB and 1 μg Trypsin was added before incubation overnight shaking at 37°C. In the morning, samples were acidified to 1% formic acid and loaded on C18 StageTips.

### LC-MS analysis

Peptides were eluted from StageTips using elution buffer (40% acetonitrile, 1% FA) and then loaded on a self-pulled 360 μm OD x 100 μm ID 20 cm column with a 7 μm tip packed with 3 μm Reprosil C18 resin (Dr. Maisch, Germany). For pull-down experiment, peptides were analyzed by nanoLC-MS in a 90 minutes gradient from 15% to 38% buffer B (for phosphopeptides 6% to 35% buffer B) at 300 nL/min using a Thermo EASY nLC 1200 system (buffer A: 0.1% acetic acid; buffer B: 0.1% acetic acid, 80% acetonitrile). Mass spectra were collected from an Orbitrap Fusion™ Lumos™ Tribrid™ Mass Spectrometer using the following settings. For MS1, Orbitrap FTMS (R = 60 000 at 200 m/z; m/z 350–1600; 7e5 target; max 20ms ion injection time); For MS2, Top Speed data-dependent acquisition with 3 second cycle time was used, HCD MS2 spectra were collected using the Orbitrap mass analyzer(R = 30 000 at 200 m/z; 31% CE; 5e4 target; max 100 ms injection time) an intensity filter was set at 2.5e4 and dynamic exclusion for 45 second.

### Data analysis

Mass spectra were searched against the UniProt human reference proteome downloaded on July 06th, 2016 using MaxQuant v1.6.2.6. Detailed MaxQuant settings: for phosphopeptide analysis, samples were set to fraction 1 and 5 for WT and mutant, respectively, to allow within-group “match between run”; for pull-down, “Label-free quantification” was turned on, but not “match between run”, no fractionation was set; Trypsin/P was selected in digestion setting. Other settings were kept as default. Protein network prediction and gene ontology analysis were performed using STRING database version 11.5 and gene ontology enrichment analysis was performed using The Gene Ontology Resource powered by PANTHER. For phosphoproteomic kinase predictions, NetworKIN analysis was performed on significantly enriched phosphosites for each mutant. Total probability was summed for each protein kinase and only the highest-ranking member of each kinase classification was used due to prediction overlap. For pie chart depiction, the top 15 kinase classification groups were pooled into categories based on recognition sequence characteristics.

### Recombinant protein purification

WT, W196R and L205R PKA catalytic (C) subunit, RIIα regulatory subunit, AKAP79 (amino acids 297-427) and PKI were produced in BL21 (DE3) pLysS E. coli cells (Novagen) and expression induced with 0.5 mM IPTG for 18 h at 18°C and purified as N-terminal His6-tag or N-terminal His6-GST tag fusion proteins by affinity chromatography and size exclusion chromatography using a HiLoad 16/600 Superdex 200 column (GE Healthcare) equilibrated in 50 mM Tris/HCl, pH 7.4, 100 mM NaCl, 1 mM DTT and 10 % (v/v) glycerol.

### In vitro Pull-down Assays for AKAP79-PKA and PKI-PKA complexes

Purified GST-AKAP79(297-417) or GST-PKI fusion proteins containing 3C protease cleavage sites were incubated for 3 hours with glutathione Sepharose beads at 4°C in the presence of 2 mM DTT, washed five times in binding buffer (50 mM Tris pH 7.4, 0.1M NaCl) and resuspended at a final protein concentration of ~5μM. C-subunit (~16 μM final concentration) and RIIα-subunits (~4 μM final concentration) were incubated with AKAP79(297-417) beads in the presence or absence of 1 mM cAMP for 20 mins at 30°C with constant agitation. The supernatant was then removed, and after three washes in binding buffer, complexes were eluted from the beads by incubation with 250 ng of 3C protease for 30 minutes at 30°C. Complexes of C-subunit and GST-PKI were isolated as above, and eluted using 1 mM GSH. Proteins were analysed by SDS-PAGE on a 12% gel.

### Protein kinase assays

PKA kinase assays were performed using a real-time mobility shift-based microfluidic system, as described previously (Byrne et al., 2016), in the presence of 2 μM of the fluorescent-tagged “Kemptide” substrate (LRRASLG) and 1 mM ATP (as standard). Pressure and voltage settings were −1.8 (PSI), −2250 V (upstream voltage), and −500 V (downstream voltage) respectively. All assays were performed in 50 mM HEPES (pH 7.4), 0.015% (v/v) Brij-35, 1mM DTT and 5 mM MgCl2, and peptide phosphorylation was detected in real time as the ratio of phosphopeptide:peptide. Changes in PKA activity in the presence of regulatory proteins (PKI and RIIα, assayed at the indicated concentrations) was quantified as the rate of phosphate incorporation into the substrate peptide (pmol phosphate · min-1 · uM enzyme-1), and then normalized with respect to control assays. To prevent ATP depletion and consequential loss of assay linearity, phosphate incorporation into the peptide was generally limited to <20%. ATP KM values were determined by nonlinear regression analysis using Graphpad Prism software. Unless otherwise specified, PKA WT and W196R mutants were assayed at a final concentration of 0.3 nM, and PKA L205R was assayed at 6 μM to account for the lower rate of activity.

### Differential Scanning Fluorimetry

Thermal-shift assays were performed using a StepOnePlus Real-Time PCR machine (Life Technologies) using Sypro-Orange dye (Invitrogen) and thermal ramping (0.3 °C in step intervals between 25 and 94°C). PKA proteins were diluted to a final concentration of 5 μM in 50 mM Tris/HCl, pH 7.4 and 100 mM NaCl in the presence or absence of the indicated concentrations of ATP, PKI or staurosporine (final DMSO concentration no higher than 4 % v/v) and were assayed as described previously (Byrne et al., 2016). Normalized data were processed using the Boltzmann equation to generate sigmoidal denaturation curves, and average Tm/ΔTm values calculated using GraphPad Prism software.

### Immunofluorescent staining

Cells were plated on acid-washed coverslips and either transfected with plasmids or infected with PKAc variant lentivirus 48 h before harvest. Cells were fixed in 4% PFA for 15 min at RT and washed 3x in PBS. Coverslips were moved to humidity chamber and blocked for 1 h at RT in 3% BSA, 0.3% TritonX100. Primary antibodies were diluted in blocking solution and applied to coverslips overnight at 4°C. Coverslips were washed 3x with PBS, incubated with fluorescent secondaries and DAPI, and washed 3x in PBS again before mounting. Images were taken on a Keyence BZ-X710 microscope and compiled and edited using Image J analysis software (FIJI).

### Pharmacology

For stress hormone measurements and acute immunoblots, drugs were applied to cells for a 10 min preincubation before washing and exchanging for fresh media and drugs for a 1 or 1.5 h incubation at 37°C. Media was then harvested and cells were lysed in RIPA (see above). For photoactivation experiments, drugs were applied to cells in glass bottom dishes ten minutes before the start of imaging. Samples use was limited to one hour after drug application to minimize variation among imaged cells.

### Mouse model

The PKA-CαR (W196R conditional knock-in) mice (Niswender et al., 2005) were made by Stanley McKnight (U Washington) and gifted to us by Diwakar Pattabiraman (Dartmouth). These mice carry a PRKACA allele encoding the W196R variant in a Cre-dependent manner. To make adrenal cortex-specific mutant mice, we crossed these to AS-Cre mice (Freedman et al. 2013), which were a gift from David Breault. Adrenal glands were dissected from acutely euthanized adult (4-7 month) mice and either snap frozen in liquid nitrogen for immunoblot analysis or frozen on dry ice for sectioning. Protein extracts were made by homogenizing glands in RIPA buffer (1% NP-40 Tergitol, 0.5% deoxycholate, 0.1% SDS, 130 mM NaCl, 20 mM NaF, 2 mM EDTA, and 20 mM Tris pH 7.5 (at 4°C) along with 1 mM AEBSF, 10 μM leupeptin, 1 mM benzamidine, and 10 mM beta glycerophosphate). All mice were cared for and euthanized in accordance with University of Washington guidelines and approved IACUC protocols.

### Human tissue

De-identified, formalin-fixed, paraffin-embedded human adrenal cortical adenoma samples were obtained from the UW Northwest Biospecimen resource. Human adrenal tissue for immunostaining was formalin fixed and paraffin embedded. Samples were deparaffinized by placing slides in 100% xylenes once for ten minutes and once for five minutes. Samples were then rehydrated by placing slides in 100% ethanol twice for ten minutes each, followed by 95% ethanol for ten minutes, 80% ethanol for ten minutes, and deionized water two times for five minutes each. Antigen retrieval was performed by placing slides in a chamber with pre-boiled 10 mM sodium citrate buffer (pH6.0). The chamber was then placed inside of a vegetable steamer for one hour. Slides were placed under cold running water for ten minutes before permeabilization in 0.4% Triton X-100/PBS for 7 min. Blocking was carried out in 5% BSA and 10% donkey serum in PBST (containing 0.05% Tween) for two hours at room temperature. For immunofluorescence, slides were incubated with primary antibodies in 5% BSA in PBST overnight at 4°C. Cells were washed 3x in PBST for ten minutes each and incubated with Alexa Fluor conjugated secondary antibodies and DAPI in 3% BSA in PBST for 1 h at room temperature. Slides were then washed six times for ten minutes each in PBST. Samples were mounted on glass slides using ProLongTM Diamond anti-fade mountant (Thermo Fisher) and cured overnight. Images were acquired using a Leica DMI6000B inverted microscope with a spinning disk confocal head (Yokagawa) and a CoolSnap HQ camera (Photometrics) controlled by MetaMorph 7.6.4 (Molecular Devices). Tissue for immunoblotting was processed using Qproteome FFPE Tissue (Qiagen 37623).

